# Unbiased choice of global clustering parameters for single-molecule localization microscopy

**DOI:** 10.1101/2021.02.22.432198

**Authors:** Pietro Verzelli, Andreas Nold, Chao Sun, Mike Heilemann, Erin M. Schuman, Tatjana Tchumatchenko

**Author notes:** Equal contribution (Pietro Verzelli, Andreas Nold).

## Abstract

Single-molecule localization microscopy resolves objects below the diffraction limit of light via sparse, stochastic detection of target molecules. Single molecules appear as clustered detection events after image reconstruction. However, identification of clusters of localizations is often complicated by the spatial proximity of target molecules and by background noise. Clustering results of existing algorithms often depend on user-generated training data or user-selected parameters, which can lead to unintentional clustering errors. Here we suggest an unbiased algorithm (FINDER) based on adaptive global parameter selection and demonstrate that the algorithm is robust to noise inclusion and target molecule density. We benchmarked FINDER against the most common density based clustering algorithms in test scenarios based on experimental datasets. We show that FINDER can keep the number of false positive inclusions low while also maintaining a low number of false negative detections in densely populated regions.

## Introduction

Super-resolution microscopy has opened up new opportunities in biological and biomedical research by providing unprecedented molecular insights into the inner workings of cells [1, 2]. Classical light microscopy can only resolve structural features that are larger than the diffraction limit of light (a few hundred nanometers)[3]. By overcoming the diffraction limit, super resolution microscopy has revealed long hidden mechanisms underlying intracellular transport processes [4] and the spatial organization of mRNA translation [5, 6]. The major feature of single-molecule localization microscopy (SMLM) is its ability to exploit the stochastic and sparse switching of fluorescence emission of specific labels binding to target molecules [7]. For example, in DNA-based point accumulation for imaging in nanoscale topography (DNA PAINT), target molecules are labelled with a short DNA strand and detected through stochastic and transient hybridization with a sequence-complementary, fluorophore labeled DNA strand [8]. Over time, each target molecule generates fluorescent detection events that cluster in space. Such clustering of single or densely packed molecules is observed for proteins in cells, e.g. the AMPA receptor in neurons [9], or on synthetic DNA origami structures [10]. In addition, SMLM data sets may contain detection events that represent ambiguous information. For instance, a super resolved image may contain false positive localizations (i.e. background noise from nonspecific fluorescent signal that does not originate from a target molecule). Furthermore, in high density regions of target molecules, localizations from multiple target molecules can overlap or form complex structures [11].

Because of the point-like nature of SMLM data, quantitative analysis opens opportunities to characterize cellular structures at the nanoscale [12]. One aim in SMLM data analysis is to group multiple detection events into a cluster, representing a single labeled protein or an assembly of densely packed proteins that cannot be spatially discriminated. Current state-of-the-art cluster detection algorithms rely on some form of prior, user-provided information. This information can be the type of localization patterns to be detected or prior experience with similar data sets. One of the most widely used [13–17] algorithms is the ’density-based spatial clustering algorithm’ (DBSCAN) [18, 19]. While DBSCAN is intuitive, simple, and fast for 2D datasets [20] the parameter choice that defines the clustering can suffer from human bias. To identify the ‘most probable’ clustering parameters based on prior knowledge, empirical methods for parameter identification have been proposed [19] (see Table S1). Another algorithm, Ordering points to identify the clustering structure (OPTICS)[21] circumvents the use of a global clustering parameter by defining borders between clusters of localizations through changes in local point density. Fundamentally different approaches to clustering have also gained traction to circumvent the shortcomings of DBSCAN [22–25]. In 2015, Rubin-Delanchy et al. introduced a Bayesian parameter finding approach [26, 27] which starts from a user generated prior parameter set for a cluster-proposing algorithm, and subsequently computes a posterior probability for each parameter set. Also in 2015, Levet et al. introduced SR Tesselation, an algorithm which segments large-scale images into polygonal regions [28, 29] and is specialized to reveal spatial structures at multiple scales [30]. Most recently in 2020, Williamson et al. developed a machine-learning approach named Cluster Analysis by Machine Learning (CAML)[31] that classifies localizations depending on their local neighborhood. This approach does not require the users to provide parameters. It also outperforms most classical algorithms in selected clustering challenges [31], but depends on training data sets.

Despite their successes, none of the current approaches have fully removed the dependency on prior knowledge – either a statistical model is needed, a reference density needs to be set, or a machine-learning model needs to be trained on a pre-selected data set. We address the problem of parameter sensitivity and user-generated bias by building on the widely applied DBSCAN [13, 32–35] algorithm and propose an unbiased parameter selection which we call FINDER. FINDER minimizes dependencies on prior knowledge by leveraging what is usually seen as a distractor: false positive localizations, or ‘noise’. The core principle of FINDER is to use information about the clustering variability with respect to the variation of the parameters to then select the most robust clustering. Global noise levels therefore act as a lower boundary for the sensitivity of the algorithm, preventing over-segmentation and minimizing false positive cluster inclusions. To validate this approach, we use clusters which we identified in super-resolution images [8] to produce synthetic test sets, and compare the performance of FINDER with one of the currently best performing clustering algorithms, the adaptive machine-learning algorithm CAML [31]. We also compare the performance to classical DBSCAN and density-based OPTICS clustering approaches [21]. We show that the FINDER algorithm is both independent of training data and is computationally tractable. FINDER also exhibits a similar or better performance as measured by true positive detections, and a reduction in false-positive cluster detections. Finally, since the parameters explored by FINDER have a precise meaning, the algorithm outcome is straightforwardly interpretable.

## Results

The assignment of clusters of localizations in super resolution microscopy is not trivial. Consider, for example, DNA PAINT data sets of super resolved neuronal AMPA receptor localizations. In Fig. 1 (left) one can see that owing to variable cluster sizes, high-noise and overlap the identification of clusters of localizations is difficult. Similarly, data sets of DNA origami trimers (Fig. 1 right) indicate that identifying localizations that belong to single molecules of interest and separating these locations from the background noise is also a challenge. Algorithms can propose candidate clusters that may correspond to single molecules of interest. However, verifying the reliability of the result is difficult because the ground truth is not always known and different parameter settings within the algorithm can lead to different outcomes. This means that the presence or absence of molecules of interest and their inferred location will vary depending on the parameter settings of the algorithm. In general, parameter settings, algorithm training and selection can become sources for clustering bias or errors.

**Figure 1:**
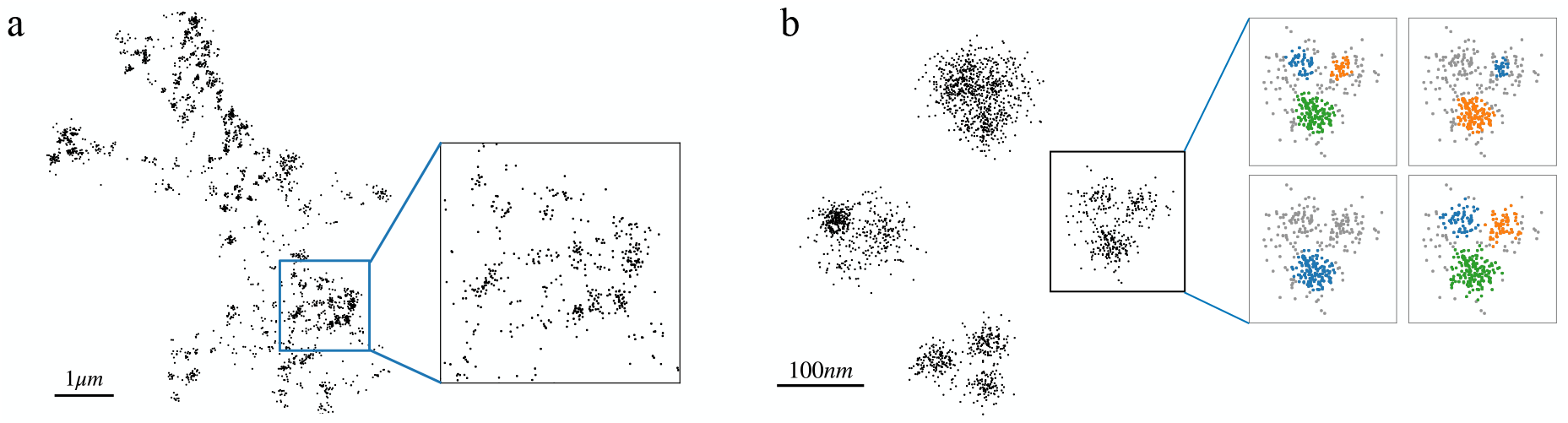
Single-molecule localization microscopy (SMLM) datasets exhibit density variations, noise inclusion and can lead to different cluster analysis results. **a** Localizations representing AMPA-receptors in a dendritic segment of a neuron [9]. **b** Example of four DNA origami trimers, with different clustering results for different parameters choices. The FINDER algorithm we propose here identifies parameters that lead to a statistically reliable assignment of clusters of localizations.

To solve these challenges we developed a new clustering approach (FINDER) which is based on a similarity metric rather than prior knowledge. We motivate the FINDER algorithm in Fig. 2 and present a step-by-step explanation in the method section “FINDER algorithm”. To benchmark FINDER, we used super resolution images of structurally well-defined DNA origami trimers and tetramers. DNA origami are folded DNA nanostructures with well-defined binding sites for fluorophores [10]. As such, the ground truth of the geometry of binding sites is known: the algorithms should identify not more than 3 or 4 subclusters for each DNA origami trimer and tetramer, respectively. Note that the subclusters of the identified oligomers exhibit a considerable heterogeneity in their number of localizations per subcluster.

**Figure 2:**
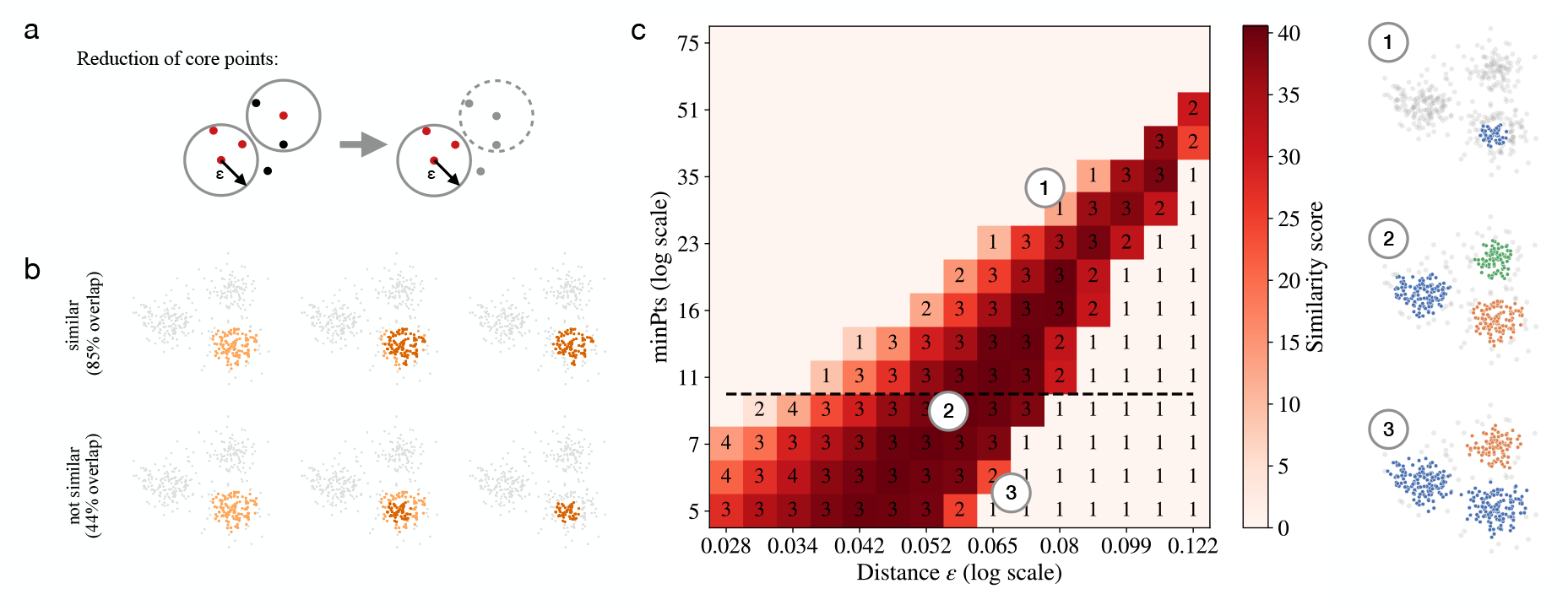
Schematics of FINDER algorithm a. In DBSCAN, a new cluster is initiated when at least one core point (shown in red) is present that has at least *minPts* other points within distance *ε* from the core point (see circles, left). Inspired by DBSCAN, the clustering algorithm used in FINDER iteratively removes non-core points (shown in black) which results in a more frequent identification of noise localizations (grey points, right). **b** Two clustering assigments are considered similar, if the number of matching localizations is greater than the number of unmatched localizations. Example of two similar cluster assignments (top row) and two non-similar cluster assignments (bottom row). **c** Phase space of possible clustering outcomes. FINDER computes a similarity score among clustering results sharing the same value *minPts* (i.e., for each line on the plot, like the one highlighted with the dashed line). (1)-(3) represent three possible clustering outcomes within the parameter space. The parameters used for (1)-(3) correspond to the location of the respective number in the phase diagram, respectively.

We start by comparing FINDER to the recently proposed machine-learning based clustering algorithms (CAML), which outperformed most classical clustering algorithms in selected clustering test cases [31]. CAML feeds density variations of the neighborhood of each point into a trained classifier to identify clusters. Here, we use the pre-trained models ’CAML 07VEJJ’ and ’CAML 87B144’ from Ref. [31], which consider the first 100 and 1000 neighboring points, respectively. In contrast, FINDER identifies an optimal parameters based on a similarity measure that is computed across an interval of probable parameters (see Methods for details). FINDER uses a density-based approach which is based on the core points defined in DBSCAN and then optimizes this parameter based on the parameter phase space in the particular data set. Core points are points that have at least *minPts* neighbors within radius *ε*. With DBSCAN, these core points are used to initiate clusters. Instead, in FINDER, all non-core points are identified as noise localizations and iteratively removed. We refer to this more conservative definition of core points as ‘noise-free DBSCAN’. Our tests suggest that this self-contained definition of core points is more robust to noise and leads to a lower number of false positive cluster detections (see Figs. S6-S11). In the supplemental information we also compare the performance of FINDER with a version of FINDER using the classical density-based algorithm DBSCAN and with OPTICS, showing a higher false positive detection rate and a lower robustness, respectively.

In Fig. 3, we show that the FINDER algorithm accurately predicts the number of binding sites of the DNA origami oligomer, even though the density of localizations is highly heterogeneous. Notably, the adaptive CAML algorithms lead to a wide variety of detected subclusters of localizations. CAML 07VEJJ detects 3-mers most often, but fails in detecting 4-mers. In constrast, CAML 87B144 is more accurate in detecting 4-mers than 3-mers. One explanation for this discrepancy is that, on average, the detected 3-mers have 156 localizations per subcluster, but 4-mers have 285 localizations per subcluster. This could explain why CAML 07VEJJ, which considers only the first 100 neighboring points, performs poorly for the tetramer dataset, but it does not explain the performance of CAML 87B144. It also does not explain the segmentation failure of CAML 87B144 if no random noise is added (see Fig. S5). This results suggest that a version of CAML which considers the first 1000 points would need to be retrained for such 3-mer and 4-mer datasets. Retraining of the model is one possibility to include global information into local clustering decisions but selecting training data while considering all aspects of the statistics to be captured can be challenging. Furthermore, the ground truth statistics regarding inter-and intra-cluster distance in experimentally recorded data sets is not *a priori* known which complicates the selection of the reference point for training data. It is therefore hard to define a good training data set without introducing user-generated training biases. This highlights a challenge that adaptive algorithms share: information about the local neighborhood is used for clustering decisions – but often, these decisions need information about the global properties of the dataset – such as noise intensity or cluster separation. Often, these global properties are assumed a priori through user-defined parameters or training data. This potential source of bias is avoided by FINDER, which systematically probes the full dataset to identify one set of global parameters for an easily interpretable density-based clustering algorithm.

**Figure 3:**
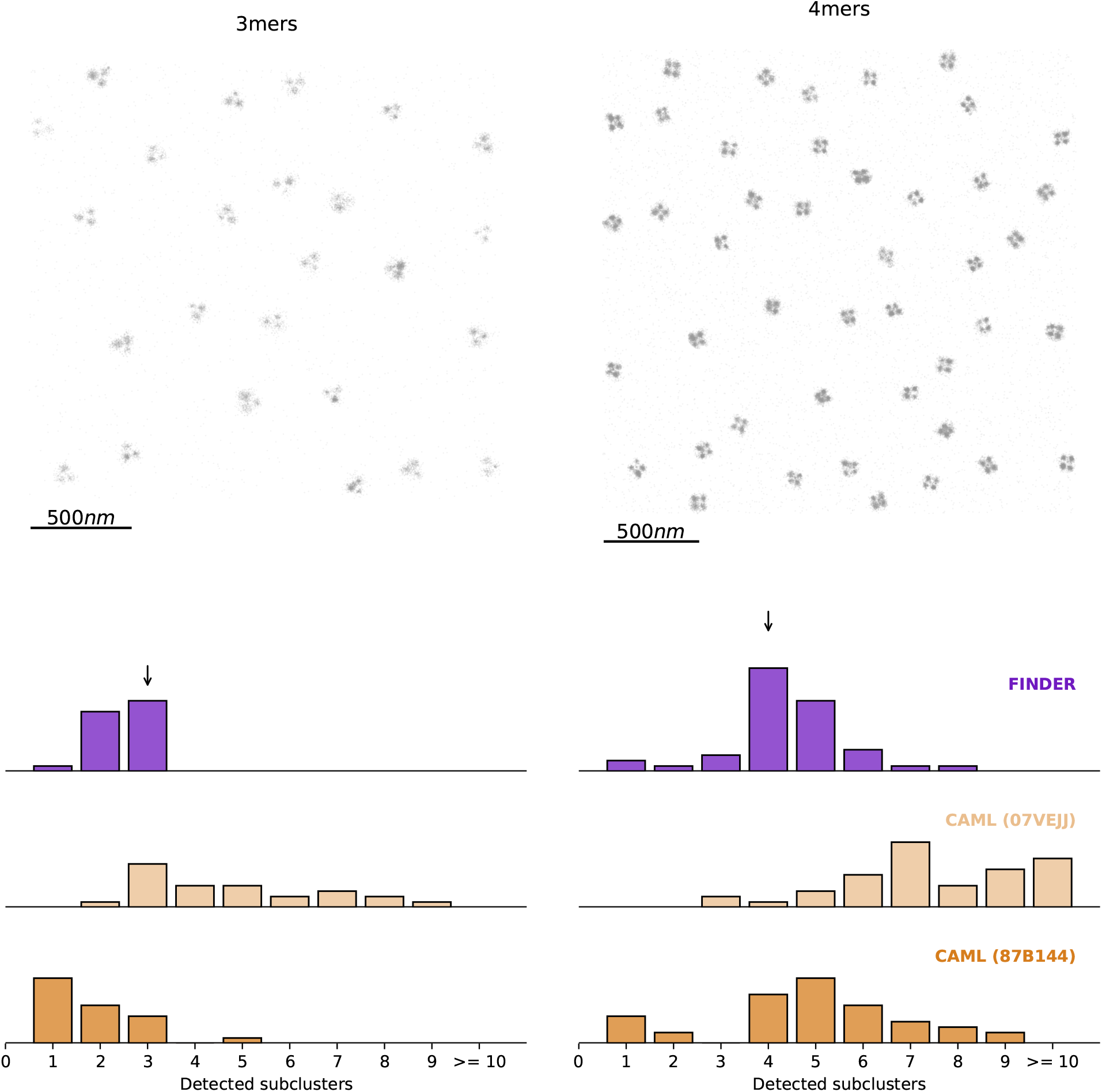
Performance of clustering algorithms for an image composed of 25 DNA-Origami 3-mers (left) and 44 4-mers (right), with added random noise localizations (10% of cluster localizations). The optimal radial parameters identified by FINDER for DBSCAN (noisefree) are *ε* = 8.05nm and *minPts* = 9 (3-mers), and *ε* = 3.61nm and *minPts* = 8 (4-mers). The histograms show the distribution of the number of subclusters detected for each 3-mer and 4-mer in our test data set, respectively. See Fig. S5 for clustering results without added noise, leading to a segmentation failure of CAML 87B144, suggesting that retraining is necessary.

As a second benchmark, we employed two libraries of unit clusters of localizations: (1) manually identified clusters in a SMLM dataset of a synapse [9], which contain on average only 17 localizations per cluster (see Fig. S1 a) and (2) manually identified sub-clusters of DNA-origami trimers, with on average 113 localizations per cluster (see Fig. S1 b). We re-arranged these unit clusters in three different configurations and added random noise points. These surrogate test data sets provide a ground-truth, while also retaining the biological variability of the cluster geometry. Our clustering outcomes are summarized in Fig 4. As expected, the CAML 07VEJJ model fails for the set of unit clusters where the number of points per cluster can be larger than the number of points considered (100, see Fig. S1 b). Interestingly, in several test cases CAML 87B144, which considers the first 1000 neighboring points, also fails for the smaller set of unit clusters (see Fig. S1 a). If the ’correct’ CAML model is chosen, performance is good, with a high number of true positive and low number of false negative detections. For all test cases we considered here, FINDER leads to a similar number of true positive cluster detections as the CAML model that performs better for the given configuration. FINDER consistently results in the lowest number of false positive cluster detections. See Figs. S6-S11 for more expansive tests. This suggests that FINDER is able to identify global parameters which give robust results with low false positive detection rates.

**Figure 4:**
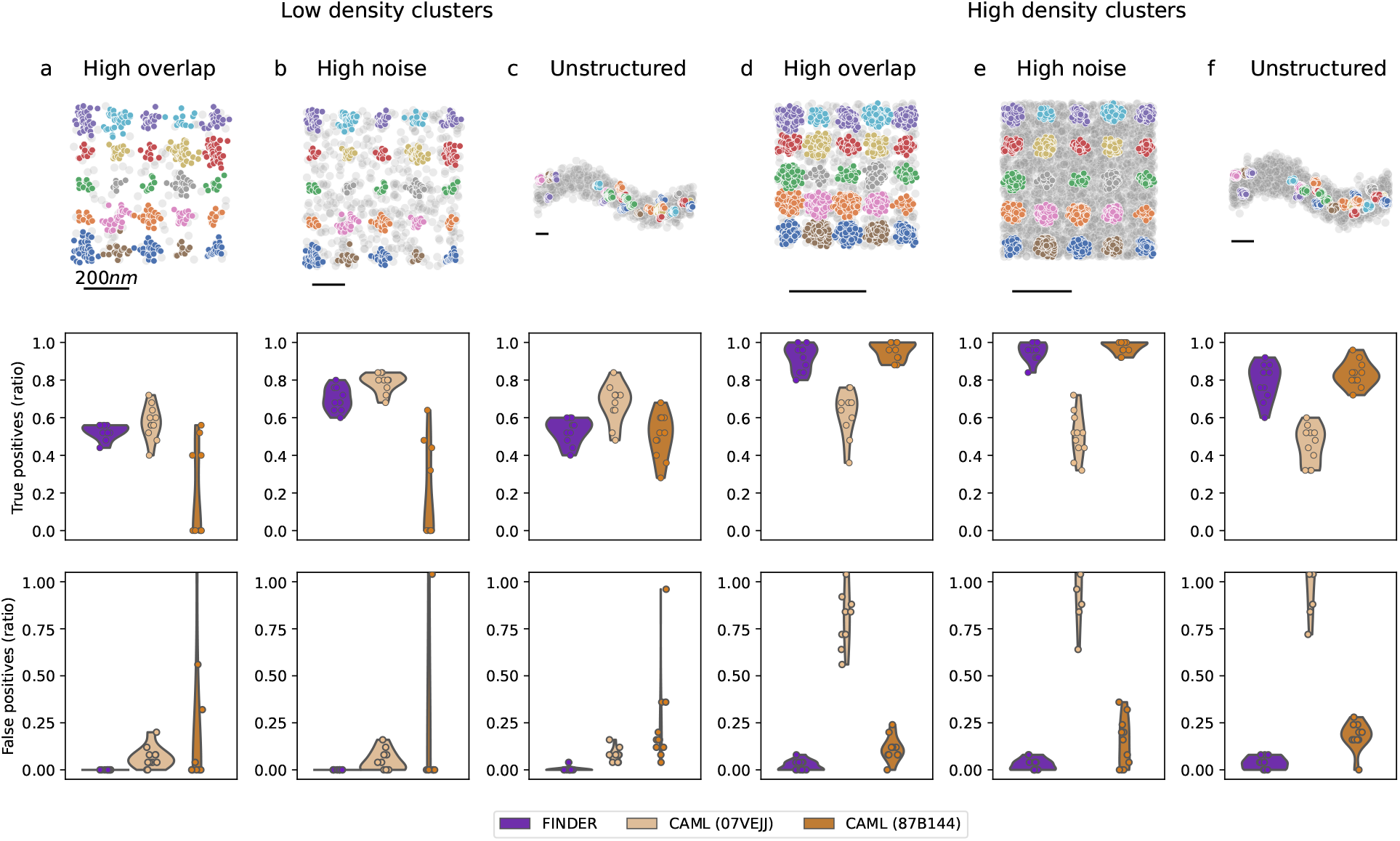
Performance of FINDER and CAML clustering algorithms across synthetic datasets which are composed of clusters of localizations from two libraries. **Low-density clusters (a-c)** are composed of clusters from a SMLM dataset of a a synapse [9] with an average of 17 localizations per cluster. **High-density clusters (e-f)** are composed of manually identified sub-clusters of DNA-origami trimers with on average 113 localizations per cluster. (a,d) **High overlap**: A grid of 5 × 5 clusters with distances equal to the maximal cluster diameter in the dataset, and with 20% random noise localizations (as a fraction of clustered localizations). (c,d) **High noise**: A grid of 5 × 5 clusters with distances equal to the 1.5 times the maximal cluster diameter in the dataset, is superimposed with an equal number (100%) of random noise localizations. (e,f) **Unstructured**: 25 clusters are randomly distributed along a sinusoidal path, with 100% and 150% added random noise localizations (as a fraction of clustered localizations) in c and f, respectively. The top row shows one instance of a randomized pattern for each case, with highlighted ground truth clusters. For further detail, see supplemental figures: S6-S11, and Fig. S16.

Finally, we applied FINDER and other clustering algorithms on SMLM datasets for which the ground truth is not known. In Fig. 5, we show the clustering results for single-molecule localization DNA-PAINT data of newly synthesized proteins after global homeostatic scaling in neuronal dendrites [6] (see Fig. S13 for an analogous analysis for a dataset of neuronal AMPA-receptors). Most clusters of localizations detected by FINDER and the two CAML algorithms have 50 or fewer localizations. CAML (07VEJJ) leads to an abrupt cutoff of cluster sizes at 100, suggesting that typical clusters can exceed that size, and therefore more than 100 neighbors need to be included in the analysis. The cluster size distribution obtained using CAML (87B144) leads to a tail in which clusters with more than 150 localizations are found, and 40% of all localizations are identified as noise. In comparison, FINDER identifies 21% of all localizations as noise localizations. FINDER therefore includes more large clusters, which usually represent molecular aggregations, than CAML (87B144). The distribution of cluster sizes for CAML (87B144) (see second bin) and visual inspection of the clusters suggests that there is an over-segmentation, which may require additional filtering. We conclude that the cluster-size distributions are overall robust with respect to the choice of the algorithm. We also note that clustering-results provided by FINDER have the advantage of being easily interpretable and reproducible: For example here, a cluster is defined if 8 core points are found within *ε* = 76.8nm, and a noise localization is defined if it does not have 8 localizations within that radius. Note that the selected value for *ε* is much larger compared to the previous cases to which FINDER was applied (Fig.3-4 and Supplemental Information related to them). This is due to the different distribution of the points in this recording, which has a larger number of points (more than 400, 000) and a different density.

**Figure 5:**
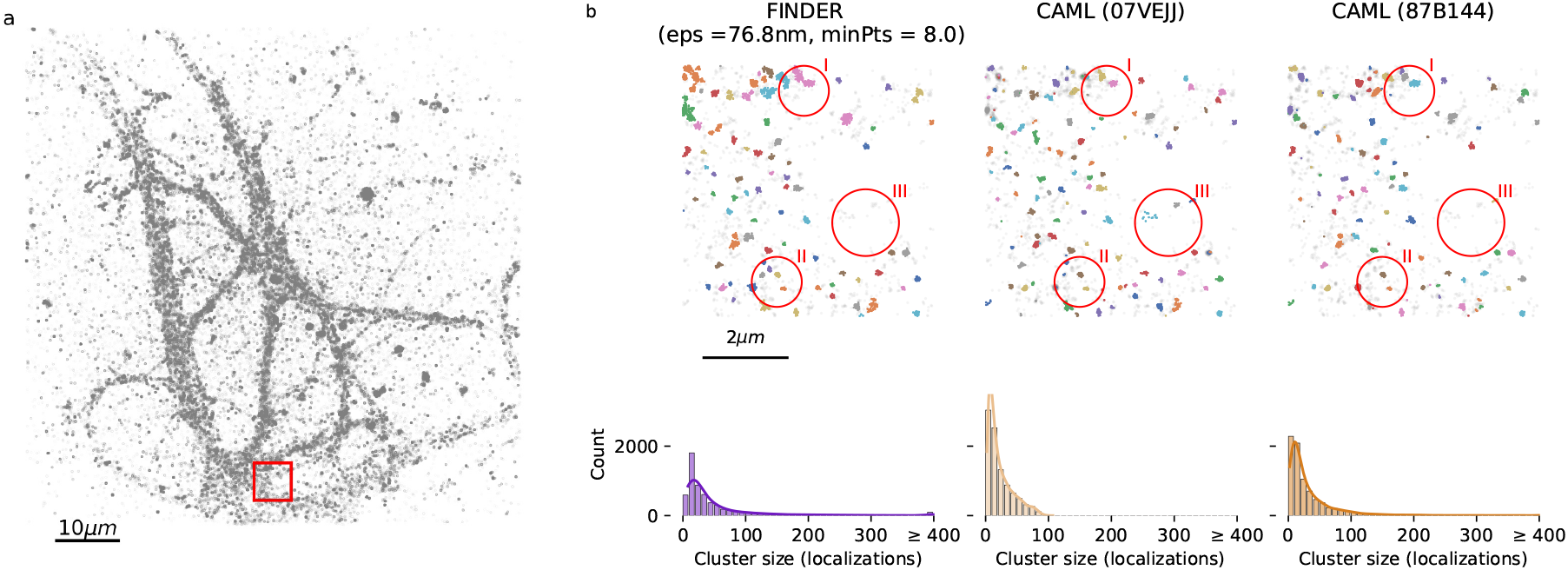
Analysis of newly synthesized proteins in neuronal dendrites in DNA-PAINT data [6]. Left: Localizations analyzed using FINDER, CAML (07VEJJ) and CAML (87B144) [31]. Right: The top row depicts a section of the full field of view corresponding to the red rectangle in the left panel. Detected clusters are highlighted as colored points and localizations that were classified as noise are shown as grey points. The optimal parameters identified by FINDER for DBSCAN (noisefree) for *minPts* = 8 are *ε* = 76.80nm. FINDER, CAML (07VEJJ) and CAML (87B144) assigned 21.8%, 0% and 1.4% of all localizations to clusters with more than 400 localizations, respectively. See red circle (I) for an example of a large cluster. The overall structure of the results is similar (eg. red circle (II)), but FINDER sets a higher threshold for the selection of small clusters. Therefore, it identifies more clusters with a low number of localizations (cluster size < 25) as noise, see eg. red circle (III). See Fig. S14 for the statistics showing the 10th-neighbor distances, Fig. S12 for an overview of the localizations not identified as noise, and for large clusters for each algorithm, and Fig. S17 for clustering outcomes within the full phasespace. See Fig. S13 for an analogous analysis of super resolved neuronal AMPA receptor localizations from [9].

## Discussion

The identification of clustered localizations in single-molecule localization microscopy data is a crucial step for quantifying proteins within nano-clusters [23, 36]. A bottleneck for automated image analysis is the identification of appropriate clustering parameters, particularly if target molecules themselves are clustered or if noise levels are high. Clustering algorithms used in scientific studies of SMLM datasets therefore need to fulfill many – sometimes contradicting – requirements. They have to be fast, robust with respect to parameter choice, and offer results that are easily interpretable. Manual parameter tuning needs to be avoided, as detailed knowledge about the system may not be readily available and can introduce human bias. Because control experiments with known ground truth clusterings are not always available, false positive cluster inclusions need to be minimized in order to avoid over- or misinterpretations of the data.

DBSCAN is one of the most popular clustering algorithms for SMLM data [13–17, 32–35]. Part of its appeal is the fact that its density-based clustering rule is intuitive and it can offer close-to-optimal results if parameters are chosen carefully [22]. It has two input parameters: a radial parameter *ε* and the minimal number of points minP ts within this radial parameter needed to initiate a cluster. However it is not always clear how to set the radial parameter manually for heterogeneous cluster sizes or datasets with changing density. Heuristic rules for setting these parameters exist [19, 22], but in many examples they are ambiguous. For instance, the optimal value for the radial parameter *ε* is commonly estimated from the k-th neighbor curves, which may be not well-defined and vary with *k* [22]. Several adaptations and improvements of DBSCAN have been proposed to deal with density-variations, and the parameter estimation problem [37, 38]. Reverse-nearest-neighbor approaches such as RECORD [39], IS-DBSCAN [40], ISB-DBSCAN [41] and RNN-DBSCAN [42] only require setting one parameter k, and can deal with density-variations within the dataset.

In many applications, however, parameters are manually chosen (see Table S1) which may not lead to optimal results when analyzing many datasets with differing localization densities.

Here, we pursue an approach that does not locally adapt to varying densities, but which sets a global threshold for the required cluster density for scenarios in which the radial parameter cannot unambiguously be extracted from k-th-neighbor curves. The idea is to leverage the information given by overall noise levels to set a global threshold for cluster inclusions and therefore minimize the number of false positive cluster inclusions. To tackle the issue of parameter choice in this scenario, while retaining the advantages of density-based clustering methods such as DBSCAN, we presented here an unbiased, automated parameter identification algorithm FINDER. FINDER combines the following benefits: First, it does not require prior knowledge about the structure of clustered localizations and as such it side-steps the need for a statistical model, the need to perform supervised training on annotated data sets, or the need for manual parameter choices that might encompass user-generated biases. Second, FINDER leads to easily interpretable results and automatically defines parameters which are then applied across the full dataset. Finally, FINDER can be applied to large, heterogeneous data sets with clustering results which are robust to noise and signal overlap (Figs. 2, 3, 4).

In the absence of a known ground truth- as is the case in most scientific exploratory data analysis-clustering algorithms need to be transparent, easily interpretable, and minimize false positive detections to avoid misinterpretation of data. In local adaptive algorithms, local density changes are employed to detect clusters of localizations. But it is often unclear on what basis the threshold for the local density change is set. In 2020, the local machine-learning based clustering method (CAML [31]) has been proposed. If the correctly trained CAML algorithm is chosen, clustering results are generally very good, with high true positive and low false positive detections (see Fig. 4). However in some cases, these methods can lead to severe over-segmentation (Figs. 3, 4, 5) and in other cases to under-counting of clusters (Fig. 3). For exploratory data analysis with an unknown ground truth, this sensitivity is critical, because one does not know which datasets are outside of the validity regime of a given trained model. This oversegmentation can be avoided if the global parameter choice is used to transfer information from the global to the local scale. For instance, information at the global scale is the overall amount of noise, or the general heterogeneity of the clusters. The local scale is the neighborhood of each localization. By avoiding pre-filtering of noise, and by performing both noise-filtering and clustering in one step, FINDER uses the global information to set the local parameters (*minPts* and *ε*). In brief, FINDER selects the parameters which leads to the most robust clustering.

We systematically benchmarked FINDER against existing algorithms using two sets of experimentally recorded clusters of localizations. We found that – despite using the same parameters across the full dataset – the cluster inclusion and exclusion criteria used in FINDER perform robustly compared to trained machinelearning models. Our tests on synthetic data sets in the high noise and the high overlap regimes showed that density variations due to noise can lead to over-segmentation in adaptive clustering algorithms if an incorrect algorithm is used (see Fig. 4). For example, for synthetic reconstructions, FINDER performed similar to the better of two pre-trained CAML models. FINDER was able to minimize the number of false positive clusters while maintaining a high ratio of true positive clusters. We also tested FINDER on a dataset of DNA-origami trimers and tetramers and showed that FINDER reduces the number of false positive cluster detections and at the same time retains many true positive detections – leading to an accurate prediction of the underlying molecular structure.

In conclusion, we showed that performing noise identification, parameter-choice and clustering in one single post-processing step, such as proposed in FINDER, provides a reliable and unbiased method for a spatial analysis of SMLM data sets. In most experimental settings, the ground truth is not known, and therefore minimizing the number of false positive cluster detections is important to avoid erroneous interpretations of experimental results. We showed here that an all-in-one cluster identification can help limit the effect of human biases, and can speed up the interpretation of single molecule microscopy datasets.

## Methods

### FINDER algorithm

The FINDER algorithm identifies the hyperparameters for a cluster-proposing algorithm. Here, we employ two cluster proposing algorithms: DBSCAN, as well as a version of DBSCAN based on iterative removal of non-core points (see Methods-section on “Noise-free DBSCAN” for details). Both algorithms take two parameters, which are a fixed minimum number of neighboring points (*minPts*), and a typical distance (*ε*). FINDER determines the optimal parameter pairs (*minPts*, ε\**) through the following steps:

1. Compute the distribution of the distance to the *k*th-nearest neighbors. Here, we set *k* = 10 (see discussion, and Fig. S17 and Fig. S18).
2. Define the interval in which the algorithm will search for the parameter *ε* as the interval between the 10th to the 90th percentile of the distribution of *k*th-nearest neighbors. The algorithm will explore *n* points linearly or logaritmically distributed in this interval. In our experiments, we set *n* = 15 and set a logaritmic scale. Here, *n* governs the numerical precision of the final parameter values and the speed of convergence.
3. The interval for the second parameter, i.e. *minPts*, is a collection of integer values. In our experiments, they span from 5 to 20. Since these are close to the biologically plausible lower and upper limits for cluster size, but other choices are also possible if they cover the biologically relevant domain.
4. For the input dataset, compute clustering results for every possible parameter combination using the cluster-proposing algorithm.
5. Compute the similarity score among clustering results sharing the same *minPts* value, see Methods section “Similarity of clusterings and similarity score”.
6. For each *minPts*, the value of *ε* that correspond to the clustering with the largest similarity score is selected. This list of pair of parameters is referred to as the *line of optima.*
7. The values for the similarity score of the elements of the line of optima are re-scaled so that they span from 0 to 1. This is accomplished by removing the minimal value from each element and then dividing by the maximum value.
8. The selected parameters are chosen moving along the line of optima. They are selected to be the first for which the normalized similarity score fall below *α* = 0.5, i.e., when its value is less than 50% of the highest similarity score.

The procedure is illustrated in Figs. S3 and S4.

### Noise-free DBSCAN cluster definition

DBSCAN initiates clusters using core points. Core points are points which have a at least *minPts* neighboring points within a distance *ε*. Here, we modified this classic DBSCAN cluster definition to make it more robust to noise. First, we iteratively remove all non-core points from the dataset of localizations *X* such that only core points remain (see Fig. S2). Next, FINDER partitions the remaining core points into clusters. The algorithm is illustrated by the following pseudo-code:

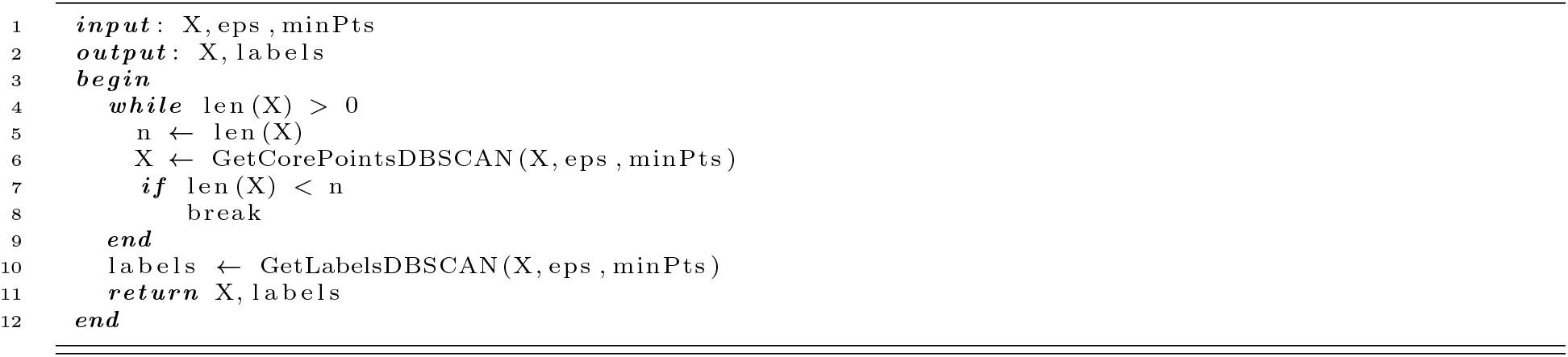

In all figures of the main manuscript, we used the noise-free DBSCAN cluster definition inside FINDER. For comparison with the classic DBSCAN cluster definition see Figs. S6-S11 in the supplementary material.

### Similarity of clusterings and the definition of the similarity score

Let 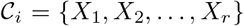 be a clustering of a set of points 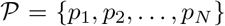. We define the similarity score of two clusterings 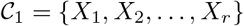 and 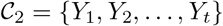 as the sum of similar subsets within the partitions:

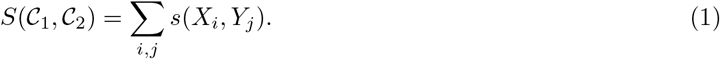

Two subsets *X_i_* and *Y_j_* are said to be similar if the number of overlapping points is larger than the number of non-overlapping points for each of the subsets:

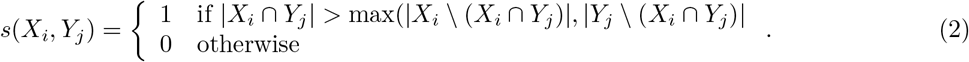

The similarity score of a clustering 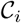 within an assembly of clusterings 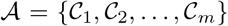 is defined as the sum of similarity scores:

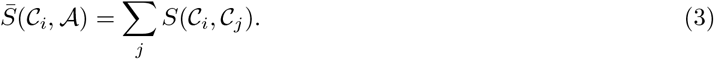

### Generation of surrogate test data from DNA origami images

To benchmark the performance of FINDER on experimental data sets and compare its performance to alternative existing clustering methods we used a DNA origami data set with a known cluster structure. We considered images of DNA origami containing three or four binding sites that we measured using DNA-PAINT [8]. Even though three or four localization clusters are expected for this DNA origami data set and this knowledge can provide a ground truth for clustering outcomes, we found that all clustering methods consistently detected a varying fraction of dimers. Upon visual inspection, we found that some expected trimers or tetramers appeared incomplete because some fluorophore binding sites were absent from some origami. We thus divided the trimer data into two groups: visibly resolved and visibly un-resolved trimers. We also introduced the group of resolved tetramers, in order to test the performance of all algorithms on a different geometric configuration See Fig. 3 for analysis of both groups.

In a second test, we accessed the robustness of the algorithms in the limit of overlapping clusters, in the high-noise limit, and with geometric constraints. We manually selected clusters representing single binding sites from resolved trimers (see Fig. 4 b) and, in a second set, we manually selected from AMPA receptor images (see Fig. S1). We then re-assembled those monomers in pre-defined grid and path geometries, and with varying levels of added random noise localizations.

### Definition of true and false positive cluster detections

In Fig. 4, a cluster *X* from the ground truth clustering 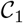 is counted as being correctly detected by cluster *Y* from clustering 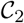 if the overlap between both clusters covers at least 30% of the points of cluster *X*, i.e. if |*X* ⋂ *Y*| > 0.3|*X*| and if cluster *Y* does not detect any other clusters of clustering 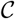, i.e. one cluster *Y* cannot detect two clusters from 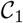. All clusters from 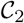 that can be attributed to a cluster of 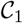 in such a way are counted as true positives, and the remaining clusters of 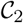 count as false positives. In Fig. S16, we vary the overlap threshold between 0% and 100%, and show robustness of the results with respect to this variation.

Note that other metrics for the similarity of two clusterings [42] such as the Adjusted Rand Index (ARI) [43] or the Normalized Mutual Information (NMI) [44], mix the similarity and the number of correctly identified clusters. In contrast, for the second benchmark in Fig. 4, we focus on how many clusters have been correctly and incorrectly identified. We therefore use a metric that uses a hard, binary, threshold for individual clusters.

### Experimentally recorded super-resolution microscopy data

#### DNA Origami data

DNA origami containing 3 (’trimers’, 3-fold symmetry, 55 nm interspacing, see Fig. S15) and 4 (’tetramers’, 4 fold symmetry, 40 nm interspacing) binding sites (containing P1 docking oligos) were imaged on the same N-STORM system (Nikon, Japan) as the above reported AMPA-receptor data: an Eclipse Ti-E inverted microscope, equipped with a Perfect Focus System (Ti-PSF) and a motorized x-y stage. Total internal reflection fluorescence (TIRF) was adjusted using a motorized TIRF illuminator in combination with a 100 x oil-immersion objective (CFL Apo TIRF, NA 1.49) with a final pixel size of 158 nm. For imaging, 647 nm excitation wavelength was used, housed in a MLC400B (Agilent) laser combiner. An optical fiber guided the laser beam to the microscope body and via a dichroic mirror (T660LPXR, Chroma) to the sample plane. Fluorescence emission was separated from excitation light via a bandpass filter (ET705/72m, Chroma) and detected by an iXon Ultra EMCCD camera (DU - 897U-CS0-23 #BV, Andor). The software NIS-Elements Ar/C (Nikon) and *μ*Manager were used to control the setup and the camera. TIRF illumination was used for super-resolution acquisitions of DNA origami data with a power of 30-40 mW, which was de- termined directly after the objective and under wide-field configuration. Time-lapse datasets with 24000 frames for DNA origami trimers and 10773 frames for DNA origami tetramers and 16 bit depth were acquired at 3.3 Hz frame rate and 5 MHz camera read-out bandwidth; pre-amplification: 3; electron multiplying gain: 50. For DNA-PAINT imaging, the imaging buffer contains P1-Atto655 (CTAGATGTAT-Atto655, Eurofins Genomics) in 500 mM NaCl, pH 7.3. The P1-Atto655 concentration was 10 nM for origami trimer and tetramer data, and 0.5 nM for AMPA receptor PAINT [9].

DNA-PAINT acquisitions were reconstructed using Picasso:Localize, a module of the Picasso software, by applying a minimal net gradient of 1500. With Picasso:Render, drift corrections were applied based on the redundant cross-correlation (RCC), with a segmentation of 1000 was applied. Drift-corrected data was filtered using Picasso:Filter. To generate the DNA origami trimer and tetramer cluster datasets for cluster-identification validation, trimer and tetramer clusters were identified by eye using Picasso:Render and manually selected with a picking diameter of 2 camera pixels.

#### Newly synthesized protein data

Newly synthesized protein data was previously reported (see Ref. [6]). In brief, cultured neuron was incubated in a growth medium containing (Tetrodotoxin) TTX for 1 hr. 15 mins before the treatment ended, the neuron was metabolically labelled with AHA. The immuno-stained neuron samples were then imaged using DNA-PAINT [8].

#### AMPA-receptor data

AMPA-receptor validation data was previously reported (see Ref. [9]). In brief, cultured neurons were stained by primary antibody against AMPA receptor GluA2 subunit before fixation and secondary antibody staining, in which the secondary antibody was modified to carry a P1 docking oligo. The immuno-stained neuron samples were then imaged using DNA-PAINT [8].

## Code Availability

The code and the data used for this project are publicly available at the following link: github.com/NoldAndreas/FINDER

## Acknowledgements

C.S. and A.N. acknowledge the support by the Add-on Fellowship for Interdisciplinary Life Science (Project number: 850027) of the Joachim-Herz Foundation. C.S. is supported by an EMBO long-term postdoctoral fellowship (EMBO ALTF 860-2018), HFSP Cross-Disciplinary Fellowship (LT000737/2019-C). M.H. is funded by the German Science Foundation, DFG GRK 2566: Interfacing Image Analysis and Molecular Life Science and DFG CRC 902: Molecular Principles of RNA-based Regulation. E.M.S. is funded by the Max Planck Society, DFG CRC 1080: Molecular and Cellular Mechanisms of Neural Homeostasis and DFG CRC 902: Molecular Principles of RNA-based Regulation and the European Research Council (ERC) under the European Union’s Horizon 2020 research and innovation programme (grant agreement No 743216). T.T. is funded by the Max Planck Society, DFG GRK 2566: Interfacing Image Analysis and Molecular Life Science and DFG CRC1080: Molecular and Cellular Mechanisms of Neural Homeostasis. We thank Ann-Christin Andres and Nina Deussner for contributing the tetramer and trimer origami data. A.N. thanks Carlos Wert-Carvajal for proof-reading and valuable feedback.

## Supplemental Information

**Table S1:** Overview of hyperparameters used for DBSCAN for the analysis of SMLM datasets. An overview of all clustering methods can be found in Ref. [23]. davg is defined as the average of the minimum distance between two points.

| Publication | SMLM method | Target molecule | minPts | r | Rationale |
| --- | --- | --- | --- | --- | --- |
| Endesfelder et al. [13] (2013) | PALM | RNA Polymerase | 4 | 30nm | *minPts* as recommended in [18] |
| Virant et al. [32] (2018) | dSTORM | various targets | 6 | 40nm | refer to Ref. [13] |
| Sanchez et al. [33] (2019) | PALM | VAR2CSA protein | 5 | 30nm | - |
| Shrivastava et al. (2019) [34] | STORM | Exogenous fibrillar Tau | 20 | 20nm | - |
| Shrivastava et al. (2020) [35] | STORM | fibrillar α-Syn polymorphs | 20 | 20nm | - |
| Mayr et al. (2020) [45] | 3D dSTORM | CD41&CD62p proteins | 3 | $2 \times \text{davg}$ | Tuned to filter noise |
| Mayr et al. (2020) [45] | 3D dSTORM | CD41&CD62p proteins | 5 | $1.5 \times \text{davg}$ | Tuned for cluster assignment |
| Harwardt et al. (2020) [16] | DNA-PAINT | MET and epidermal growth factor receptor (EGFR) | 10,15 | 10-15nm | nearest neighbor-based analysis (NeNA) |

**Figure S1:**
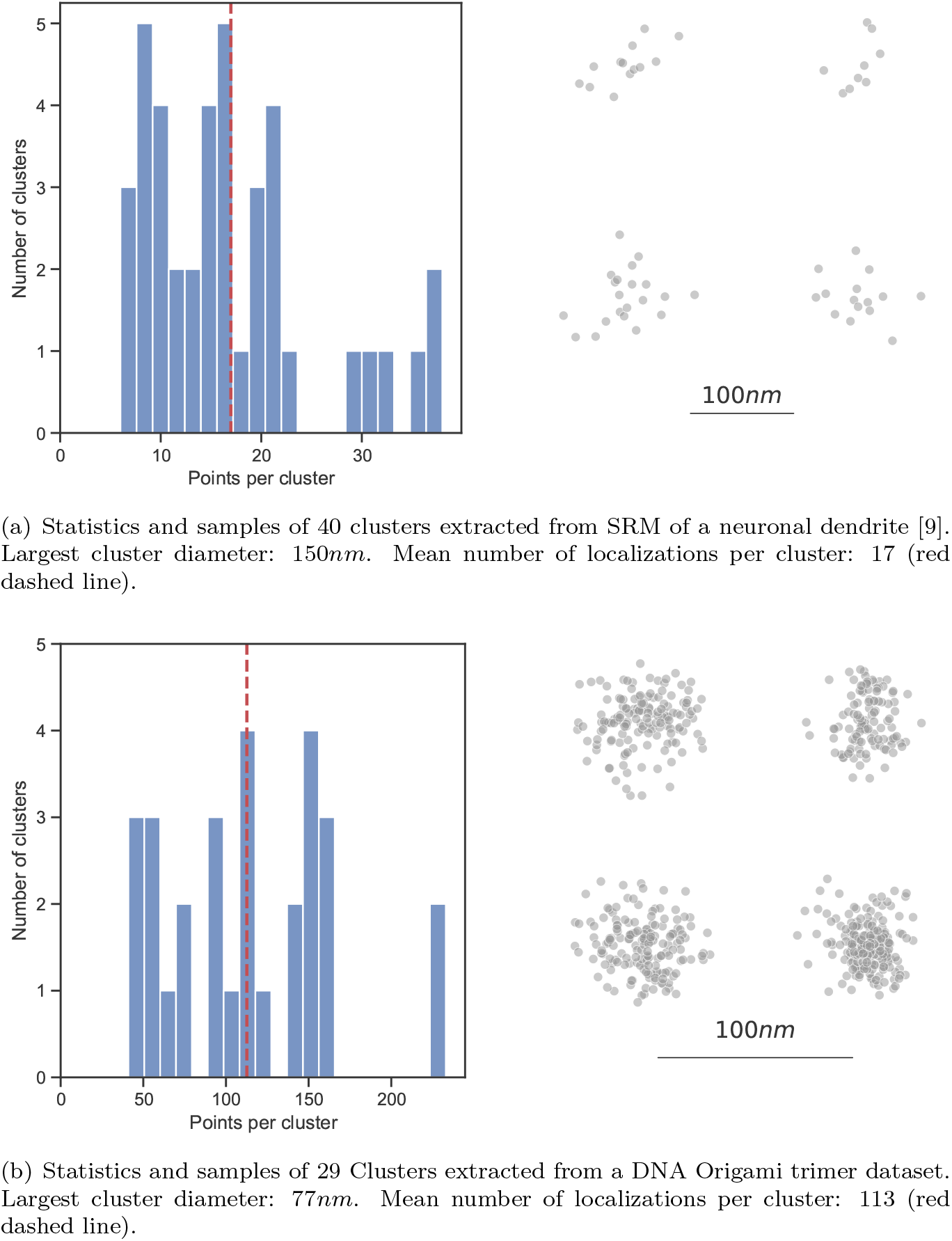
Cluster libraries: Size distribution (left) and four samples (right) of two libraries of clusters used in this work.

**Figure S2:**
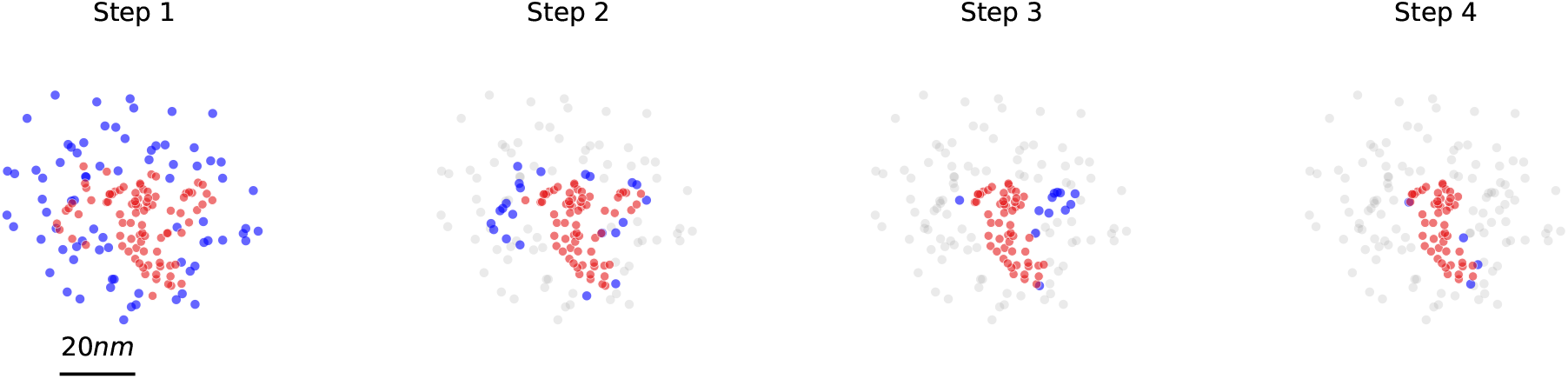
Example of the first four iterative steps of noise removal in DBSCAN (noise free). Points that have the minimum number of neighboring points *minPts* = 10 within a given distance *ε* = 7*nm* are considered as core points (shown in red). Iteratively, all points which do not satisfy this condition are removed (shown in blue), therefore restricting the number of spurious inclusions. Points that have been excluded in previous iterations are shown in grey.

**Figure S3:**
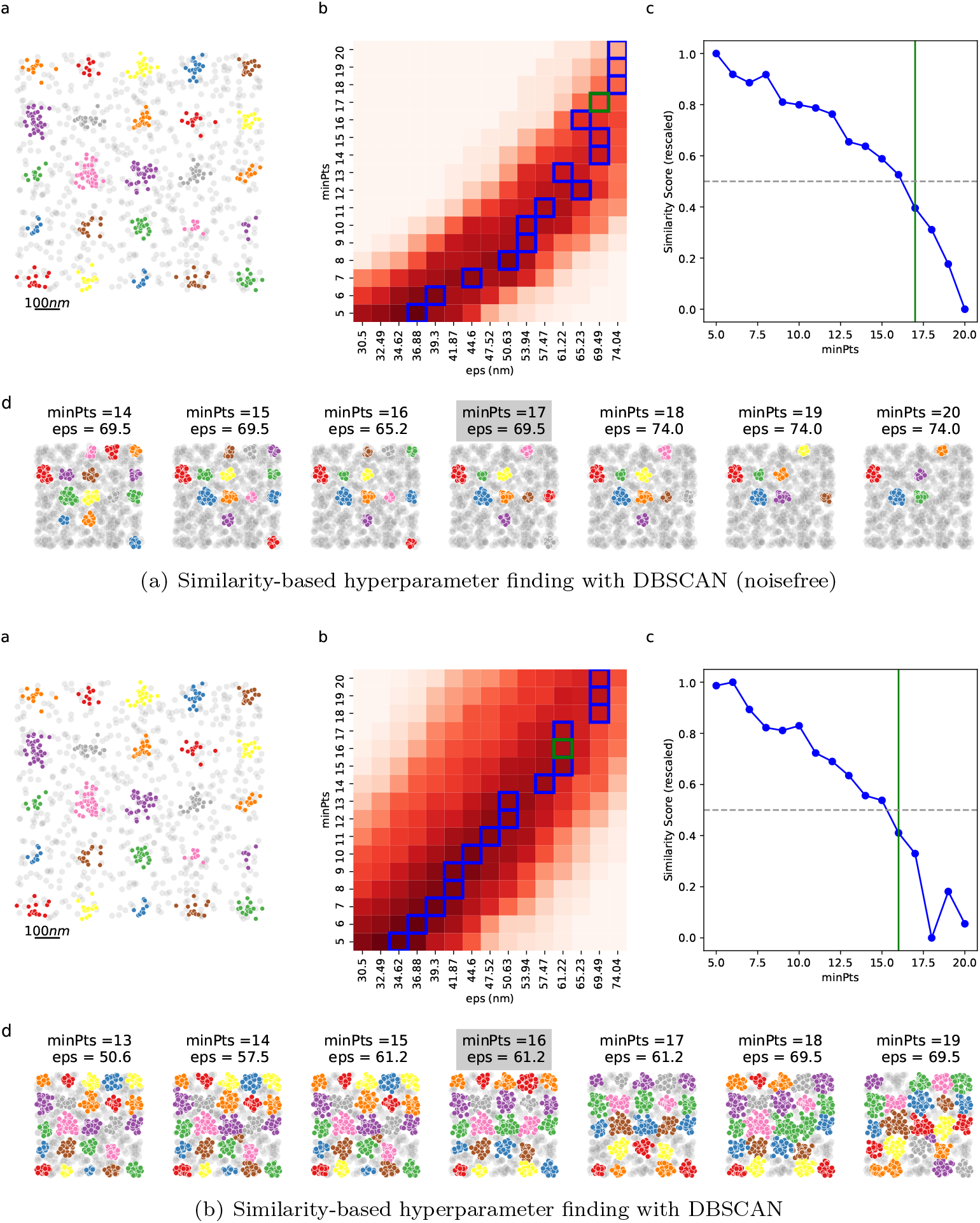
FINDER applied to configuration shown in Fig. 4(a) of the main manuscript. **Top**: DBSCAN (noisefree). **Bottom**: DBSCAN. **a**:Ground-truth clusters are shown in color, and random noise localizations in grey. **b**:The similarity score as a function of the parameters. See “FINDER algorithm” in “Methods” for a description. A darker red indicates an higher value of similarity. The blue squares highlights the points selected as the *line of optima* (step 6 of the “FINDER algorithm”). The green squares indicates parameters selected by FINDER. **c**:The parameters selection. As described in step 7 of “FINDER algorithm”, the final parameter configuration (green vertical line) on the *line of optima* is selected as the first one to fall under the threshold *α* (dashed horizontal line). Here *α* = 0.5. **d**:Clustering results for the optimal value of epsilon (grey background), and the three lower and higher values within the *line of optima*. See Fig. S19 for an analysis of the variation of the number of *ε*-values within the interval of interest.

**Figure S4:**
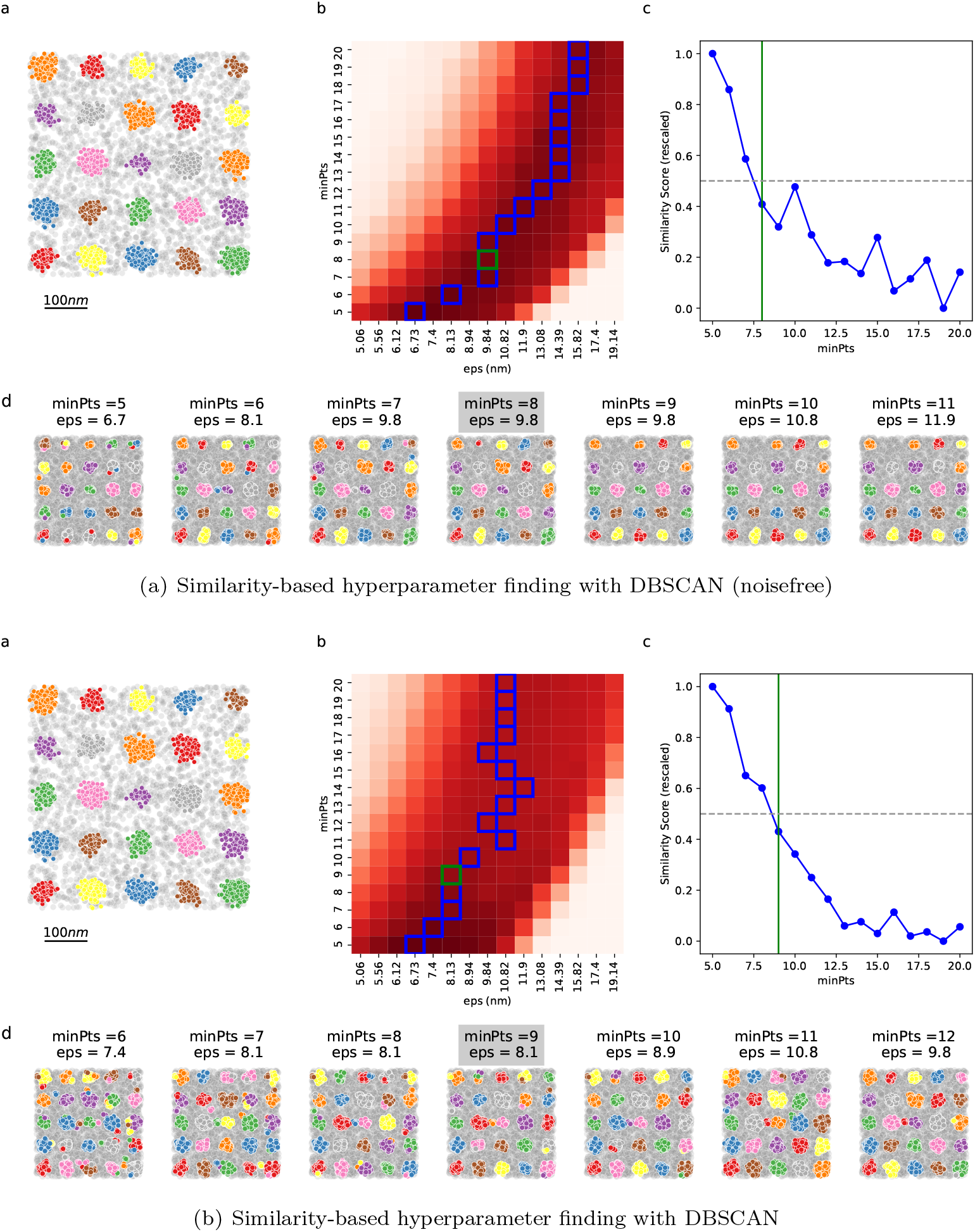
FINDER applied to configuration shown in Fig. 4(d) of the main manuscript. For details, see caption of Fig. S3. See Fig. S19 for an analysis of the variation of the number of *ε*-values within the interval of interest.

**Figure S5:**
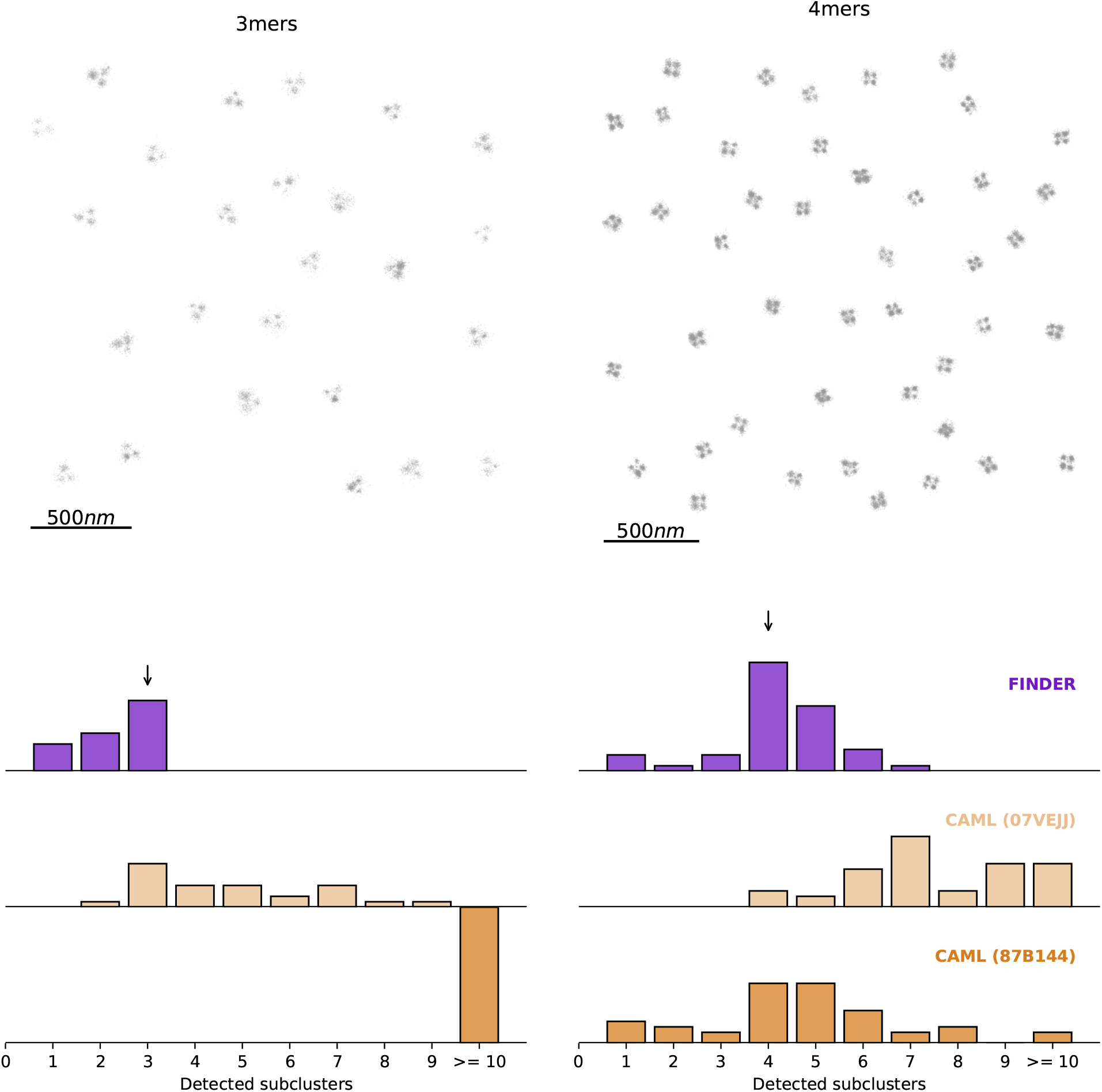
Relates to Fig. 3. Same computation as in Fig. 3, but without added random noise. Here, the optimal radial parameters identified by FINDER for DBSCAN (noisefree) are *ε* = 8.99 and *minPts* = 10 (trimer), and *ε* = 3.83 and *minPts* = 8 (tetramer). 3-mers are not recognized by CAML (87B144).

**Figure S6:**
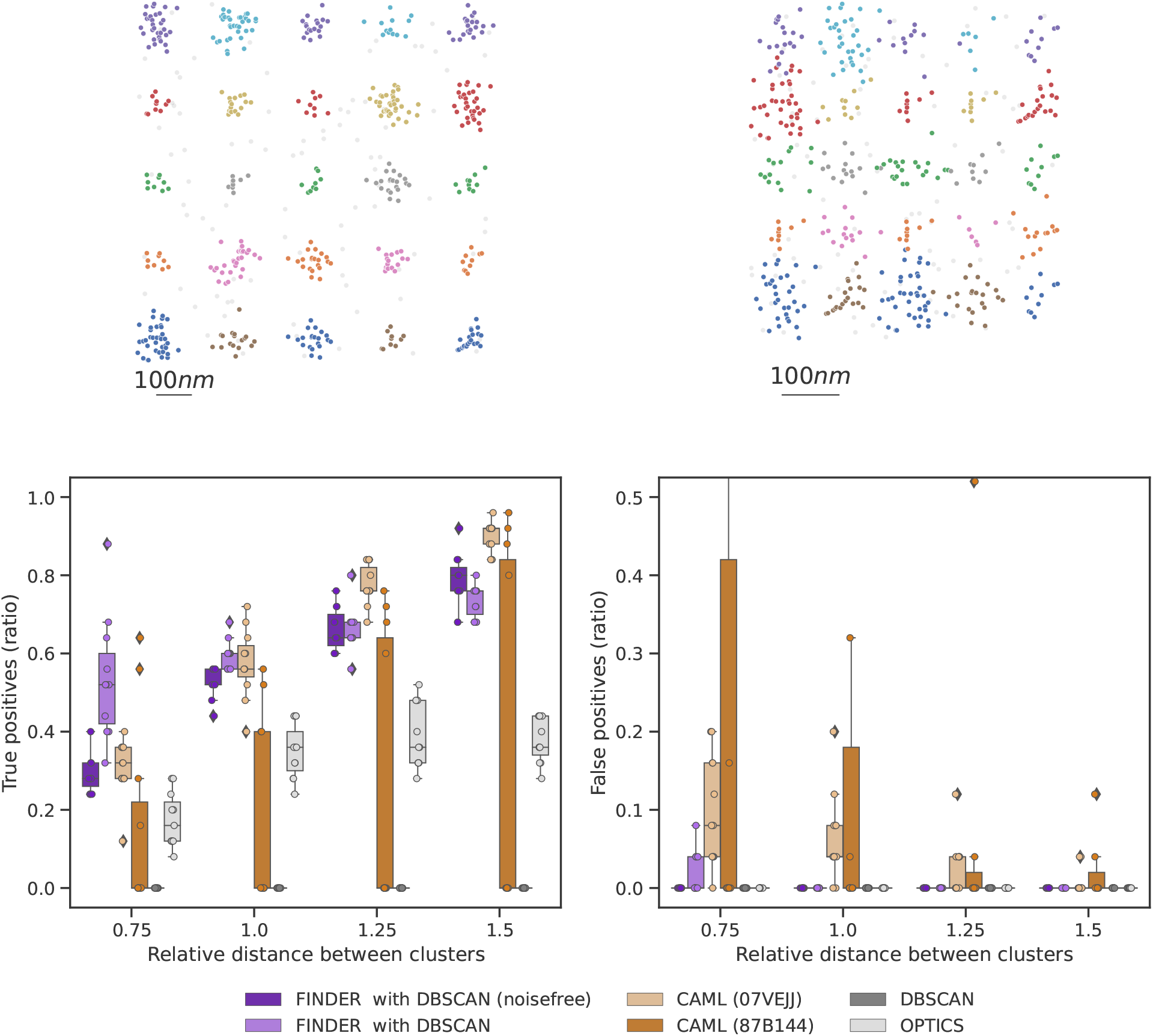
Relates to Fig. 4 a of the main manuscript. Clusters from a SMLM dataset of a synapse [9] (see Fig. 1(a)) are randomly assigned to a 5 × 5 grid. Based on the number of localizations, a ratio of 0.2 of random noise localizations was then added to the domain. The spacing in between clusters was decreased, here shown as multiples of the maximal cluster diameter of the cluster library, here 150*nm*. Top row: Two samples with highest and lowest spacing. Boxplots show the number of true and false positive cluster detections. Parameters used for DBSCAN [18] were *ε* = 10nm, minPts = 10 and for OPTICS[21]: minPts = 20, xi = 0.05, max epsilon = 100 nm.

**Figure S7:**
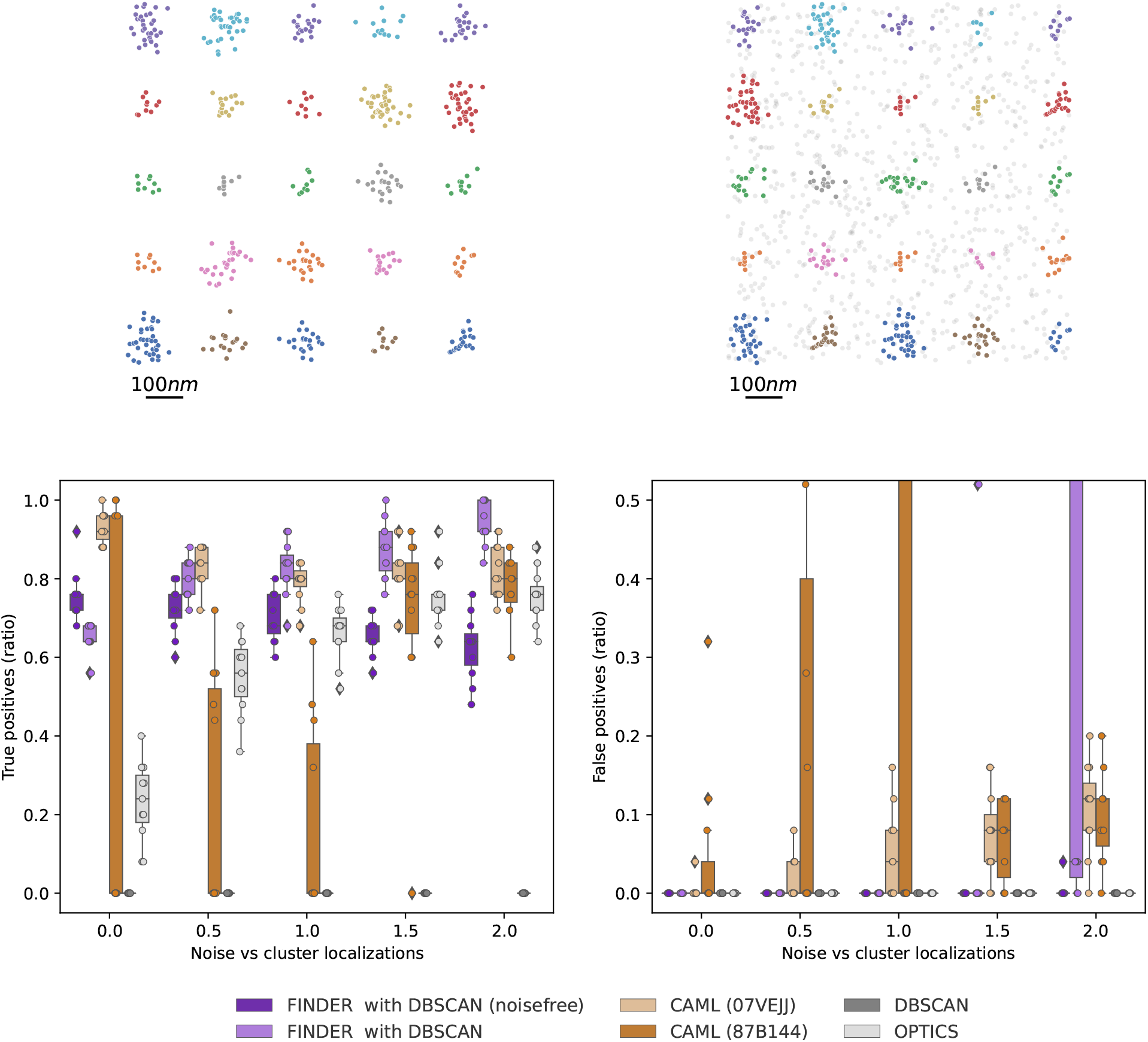
Relates to Fig. 4 b of the main manuscript. Clusters from a SMLM dataset of a synapse [9] (see Fig. 1(a)) are randomly assigned to a 5 × 5 grid, with spacing 225*μ*m. Based on the number of localizations, an increasing ratio of random noise localizations was then added to the domain. See caption of Fig. S6 for details.

**Figure S8:**
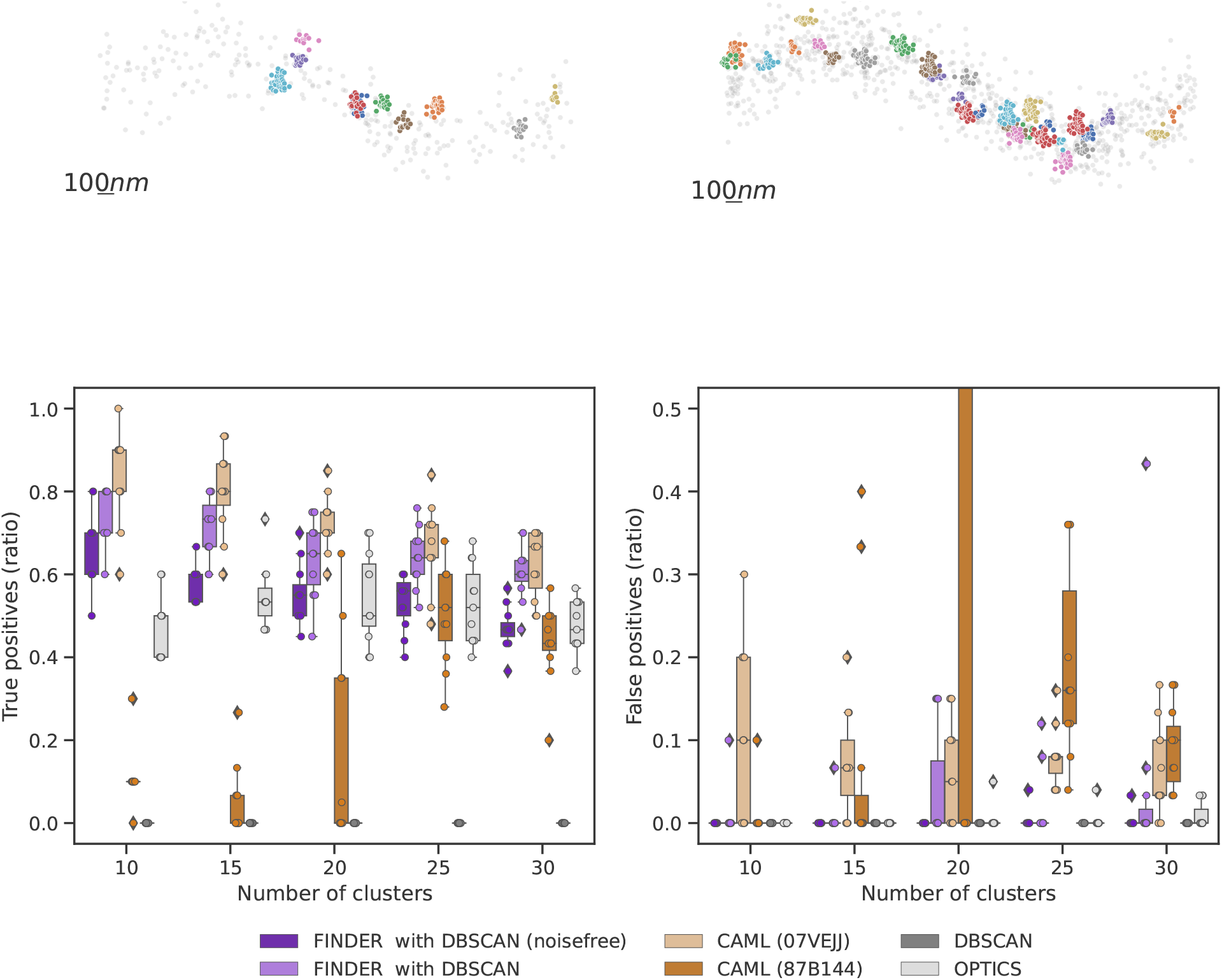
Relates to Fig. 4 c of the main manuscript. Clusters from a SMLM dataset of a synapse [9] (see Fig. 1(a)) are randomly assigned to a sinusoidal path. Based on the number of localizations of the added clusters, a ratio of 1.5 of random noise localizations was then added to the domain. The number of clusters added to the domain was then increased from 10 to 30, therefore increasing noise and overlap. See caption of Fig. S6 for details.

**Figure S9:**
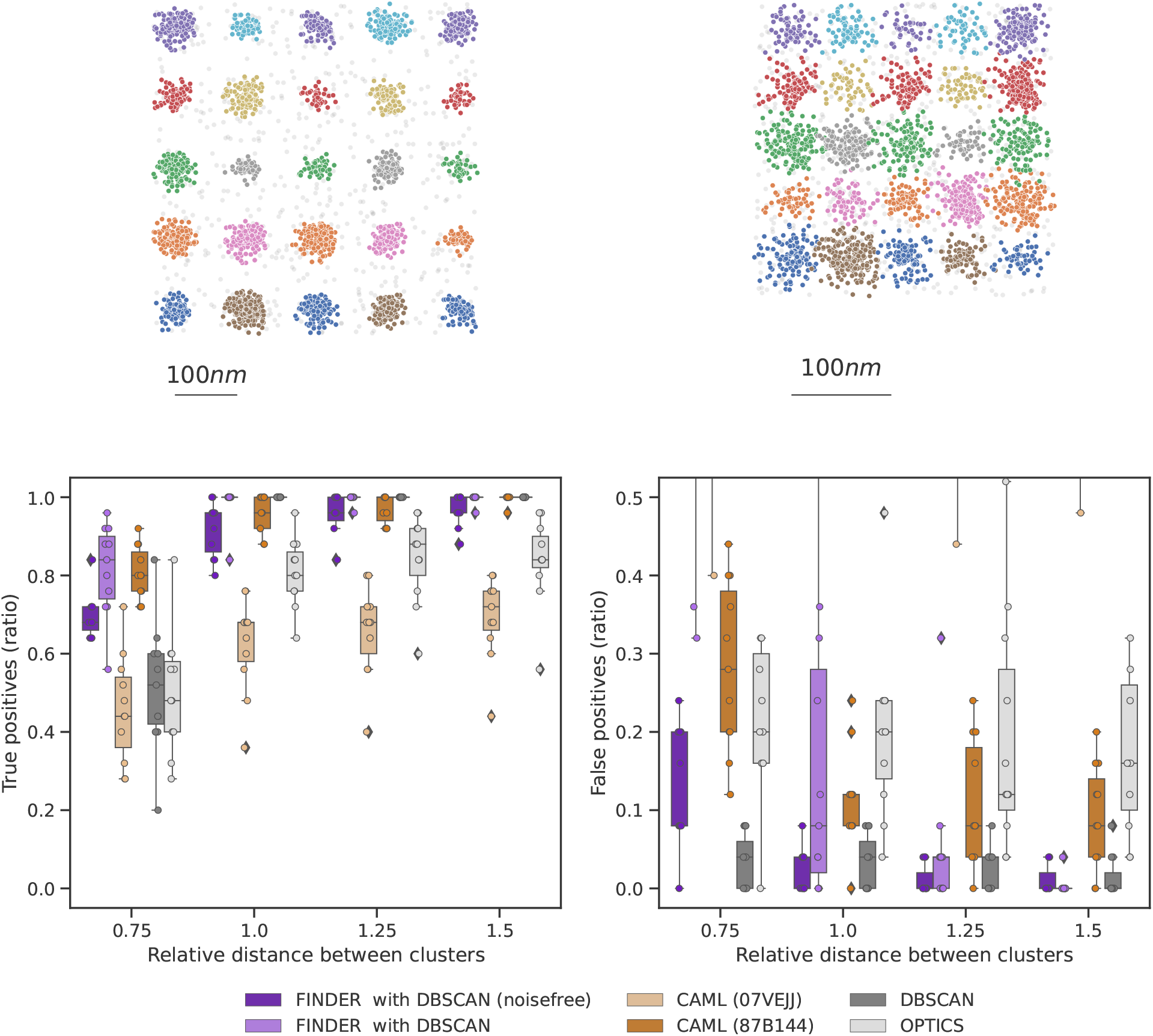
Relates to Fig. 4 d of the main manuscript. Clusters from a DNA origami dataset (see Fig. 1(b)) are randomly assigned to a 5 × 5 grid, with spacing 115*μ*m. Based on the number of localizations, an increasing ratio of random noise localizations was then added to the domain. See caption of Fig. S6 for details.

**Figure S10:**
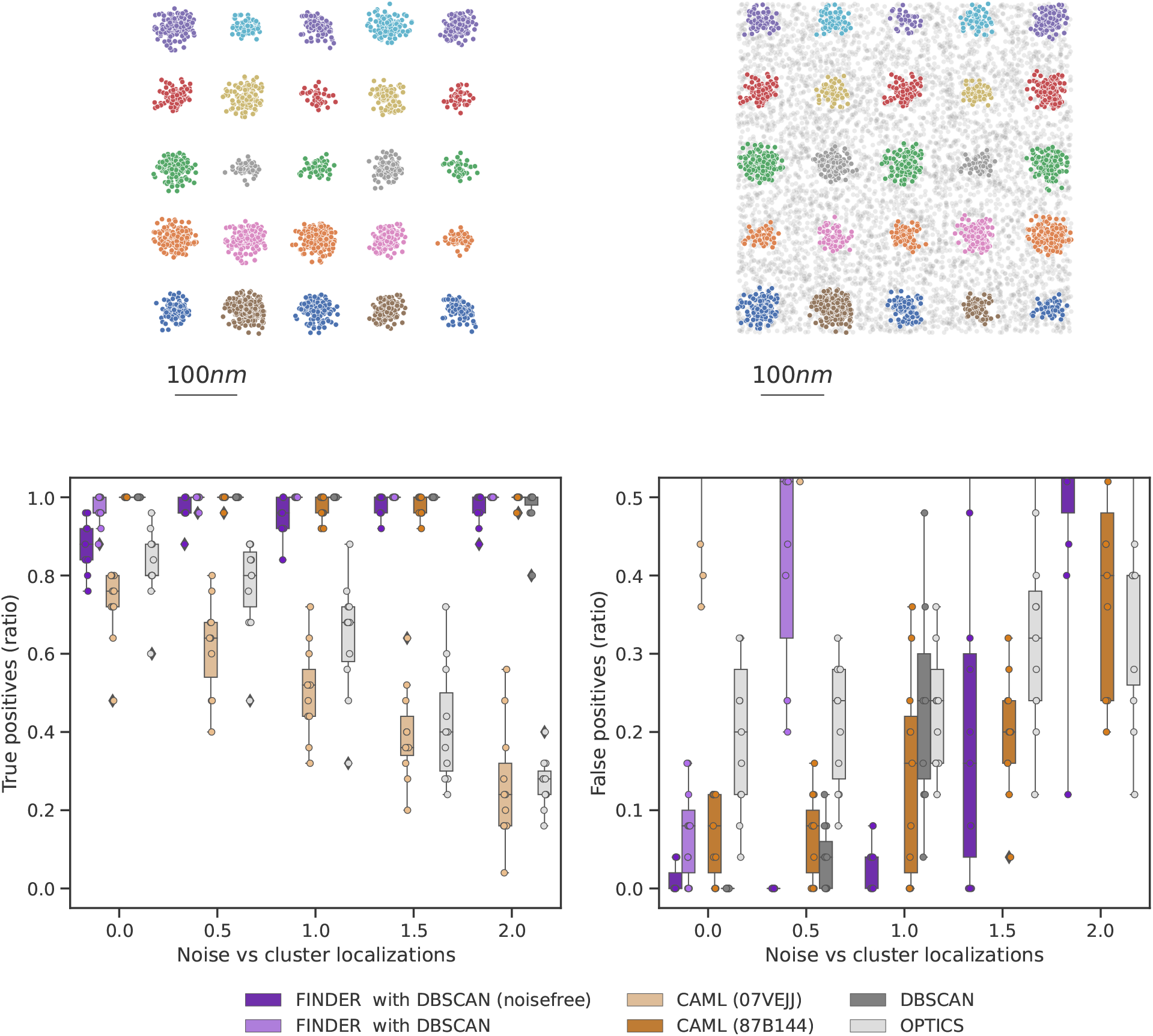
Relates to Fig. 4 e of the main manuscript. Clusters from a DNA origami dataset (see Fig. 1(b)) are randomly assigned to a 5 × 5 grid. Based on the number of localizations, a ratio of 0.2 of random noise localizations was then added to the domain. The spacing in between clusters was decreased, here shown as multiples of the maximal cluster diameter of the cluster library, here 77*nm*. See caption of Fig. S6 for details.

**Figure S11:**
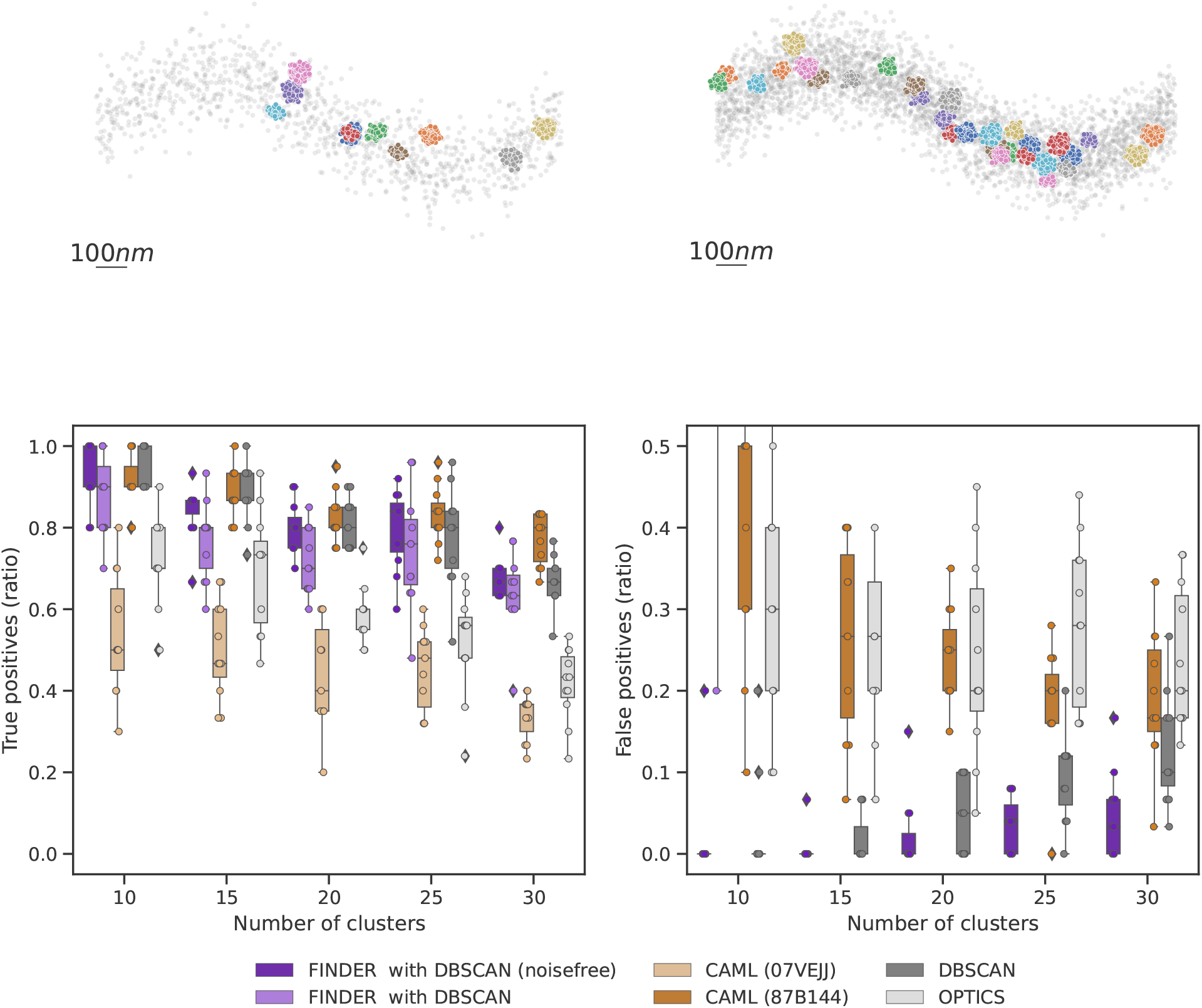
Relates to Fig. 4 f of the main manuscript. Clusters from a DNA origami dataset (see Fig. 1(b)) are randomly assigned to a sinusoidal path. Based on the number of localizations of the added clusters, a ratio of 1.0 of random noise localizations was then added to the domain. The number of clusters added to the domain was then increased from 10 to 30, therefore increasing noise and overlap. See caption of Fig. S6 for details.

**Figure S12:**
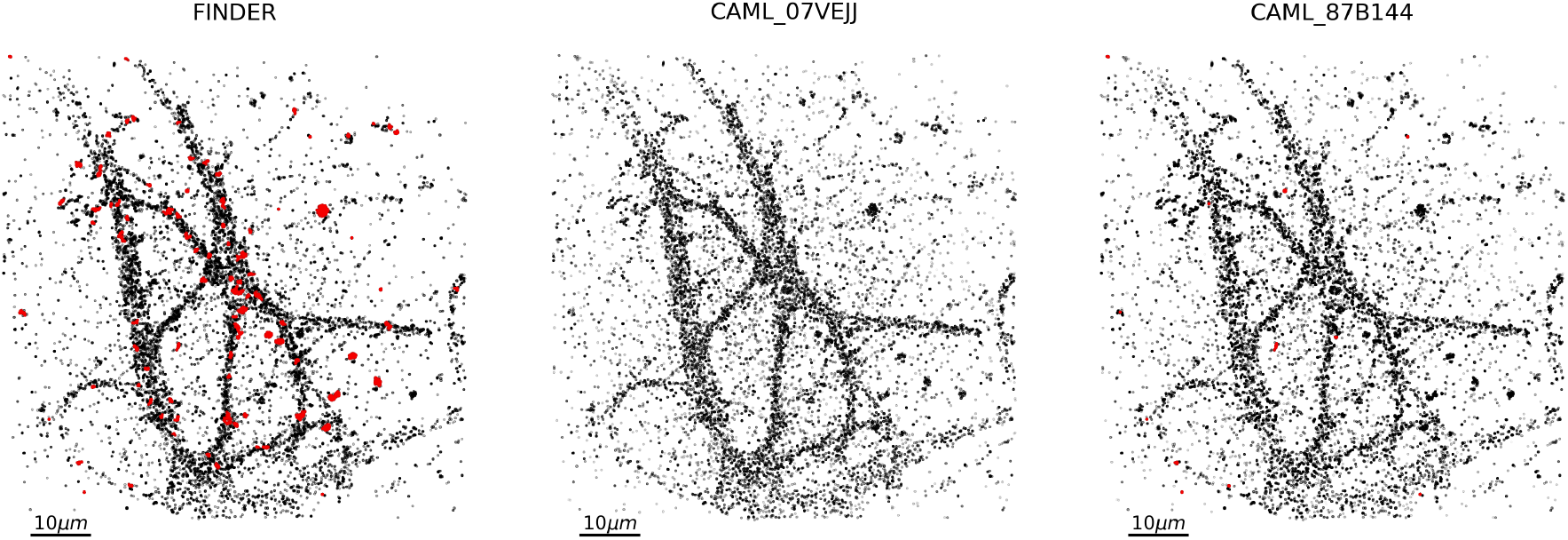
Related to Fig. 5. All localizations not identified as noise. Localizations assigned to clusters of more than 400 are highlighted in red.

**Figure S13:**
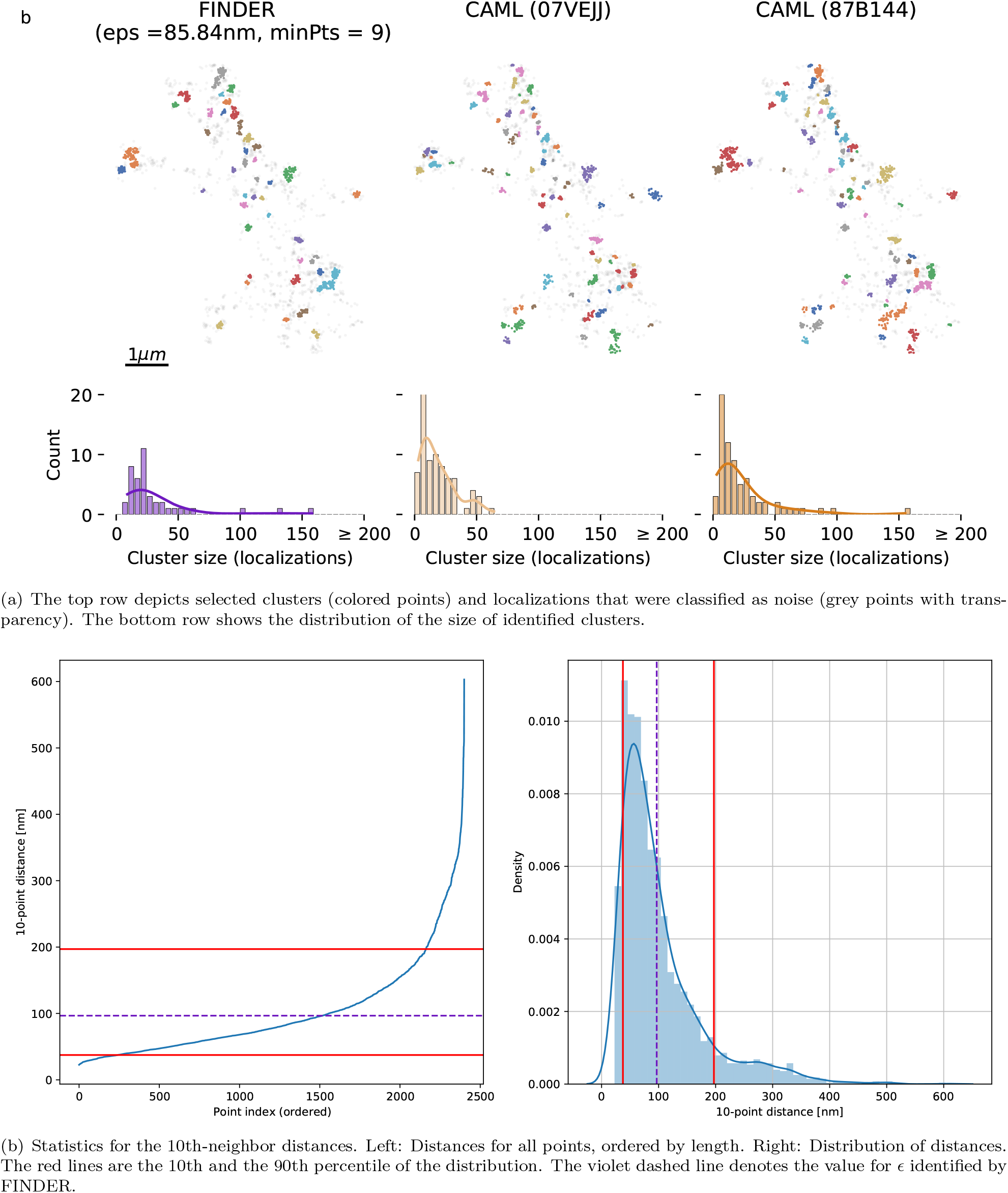
Analysis analogous to Fig. 5, but for super resolved neuronal AMPA receptor localizations [9]. As in Fig. 5, FINDER identifies fewer clusters with few localizations (cluster size < 25). See Fig. S18 for an analysis of the full phasespace.

**Figure S14:**
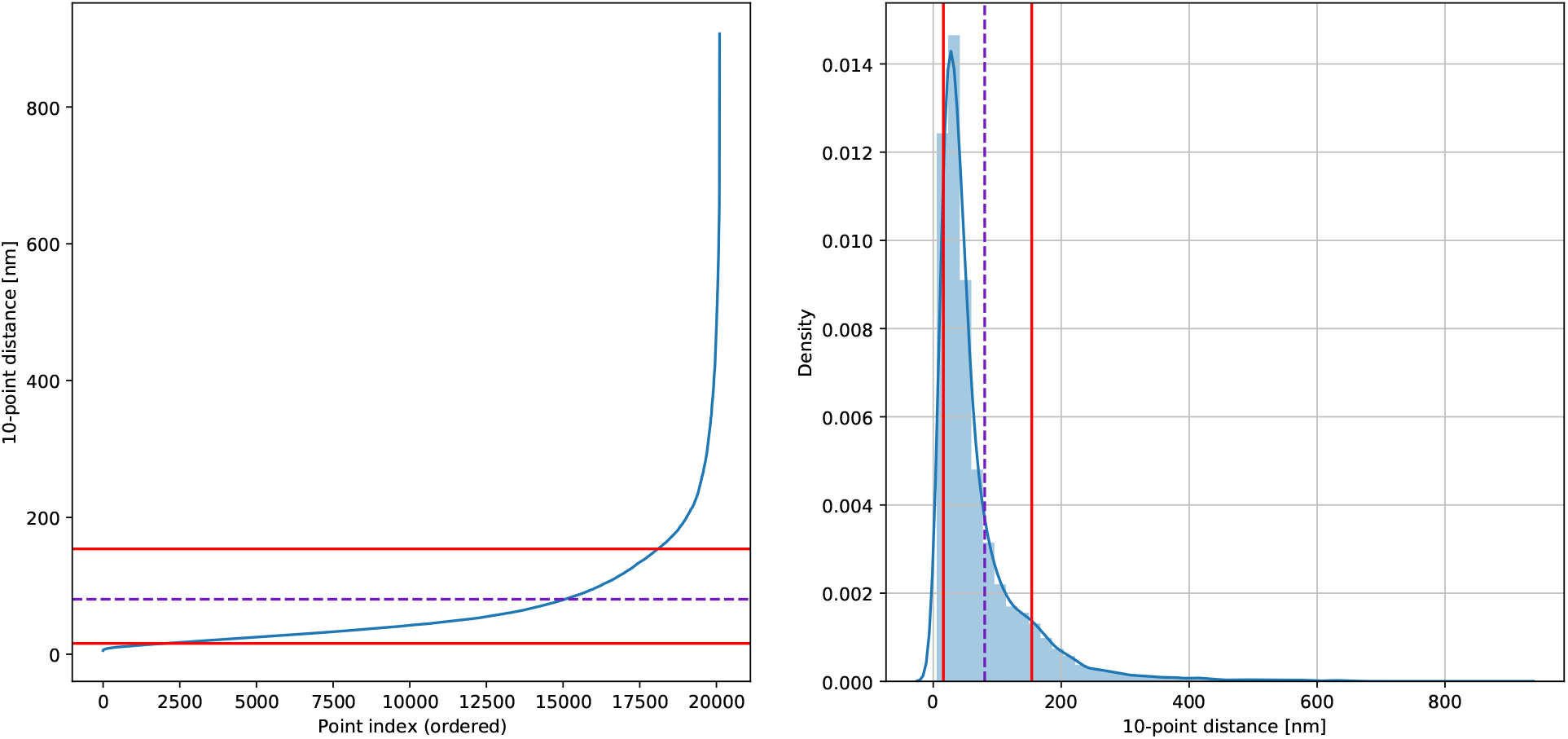
Statistics for the 10th-neighbor distances for the case analyzed in Fig. 5. Left: Distances for all points, ordered by length. Right: Distribution of distances. The red lines are the 10th and the 90th percentile of the distribution. The violet dashed line denotes the optimal value for *ϵ* identified by FINDER.

**Figure S15:**
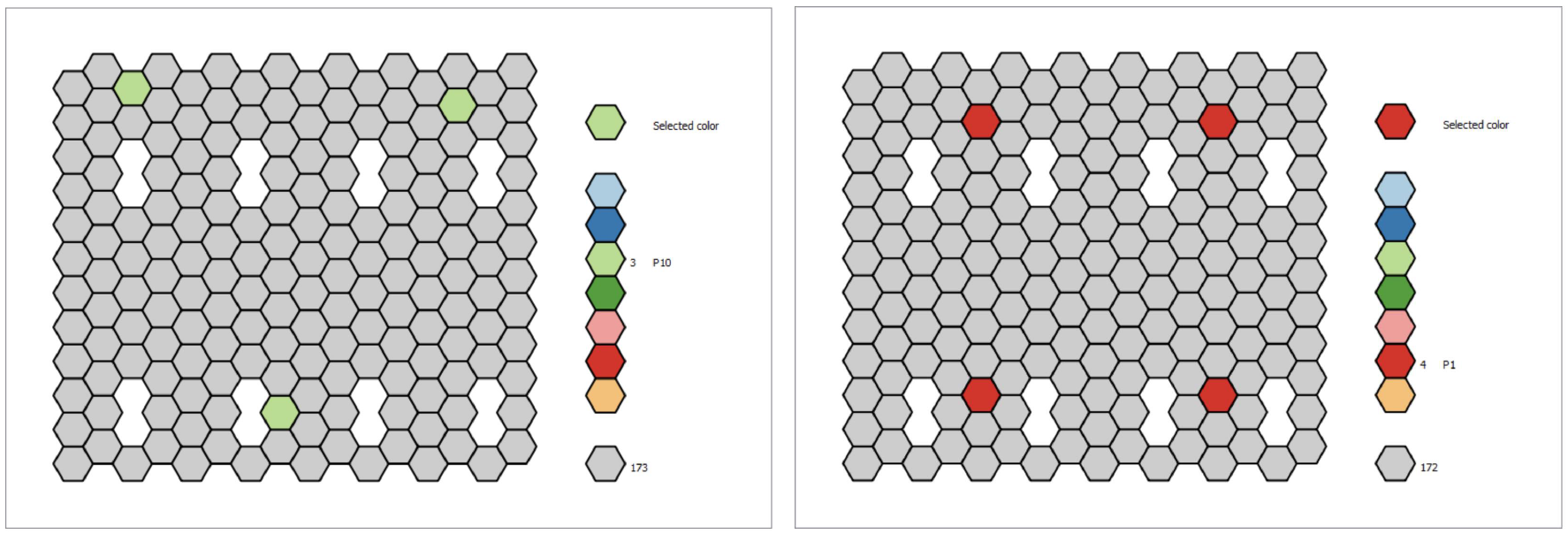
Left: 3-fold symmetric DNA origami containing 3 binding sites. Right: 4-fold symmetric DNA origami containing 4 binding sites.

**Figure S16:**
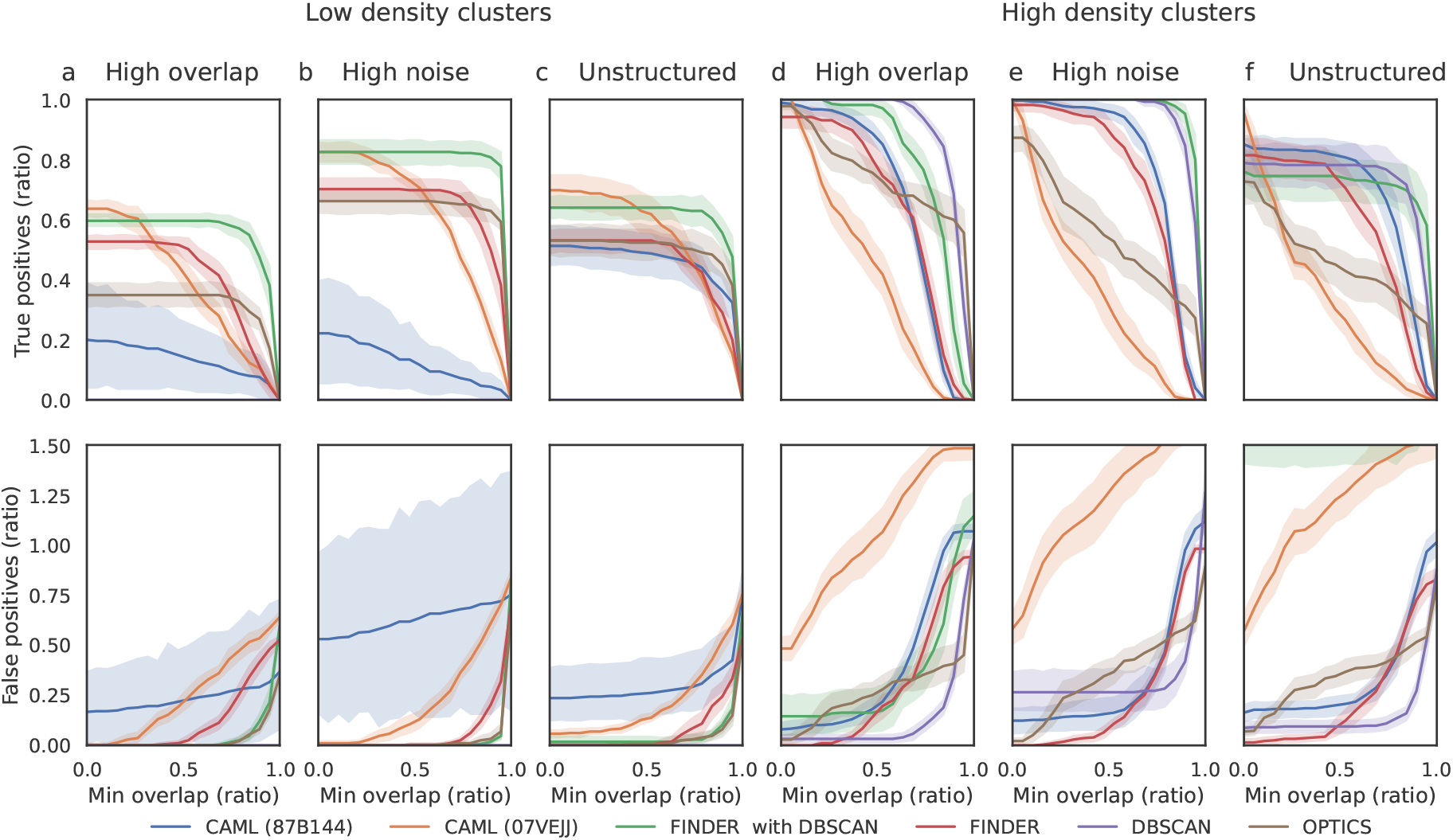
Re-evaluation of the results from Fig. 4. When comparing the clustering outcome to the ground truth, the required overlap between the detected and the ground truth cluster is varied here from 0 to 1 (100%). Note that the clustering outcome does not change with the variation of this threshold – only the ratio of clusters that are attributed to being true or false positives changes. Ratios are given with respect to the number of ground truth clusters.

**Figure S17:**
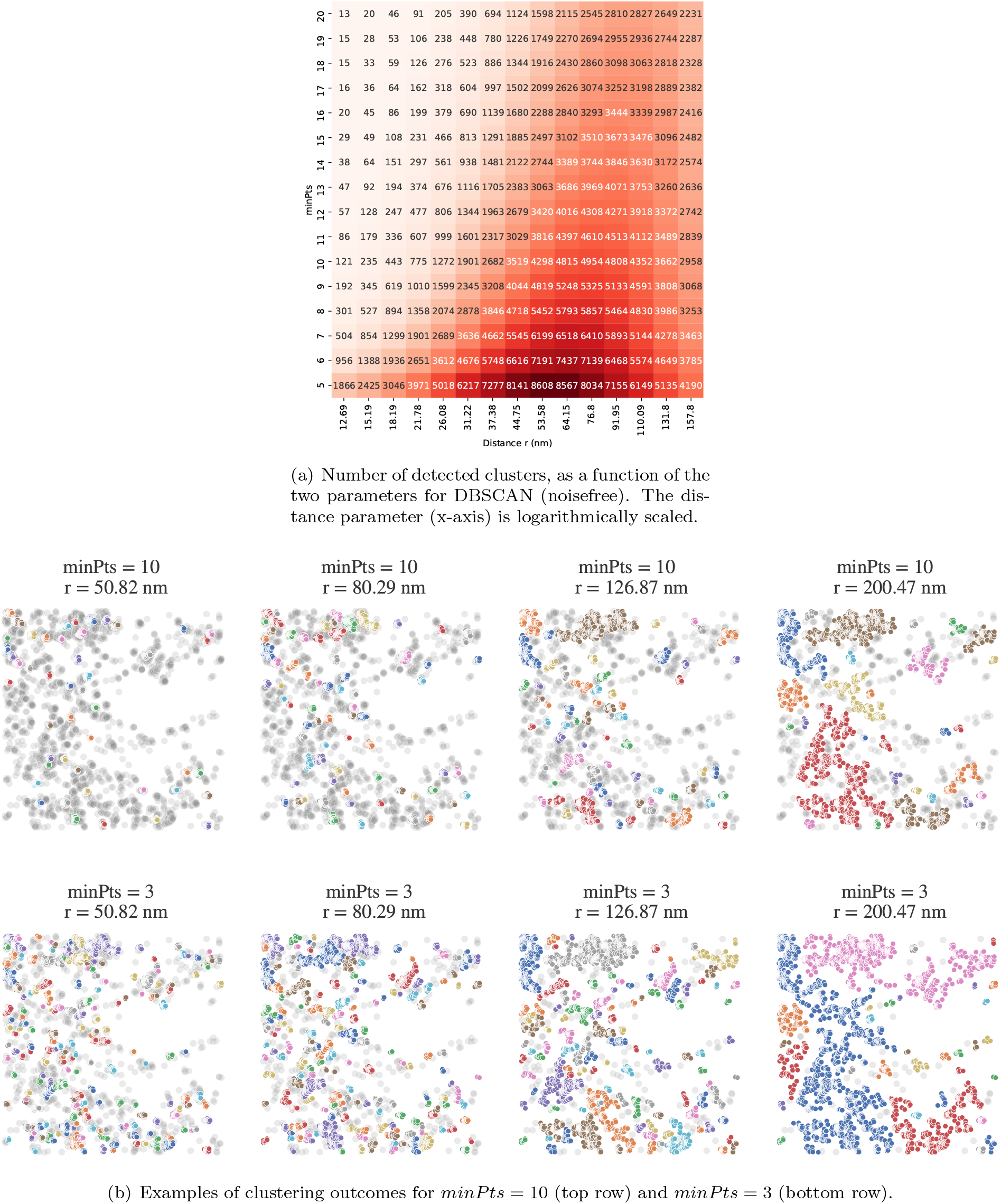
Exploration of clustering outcomes with DBSCAN (noisefree) for the localizations within the red rectangle in Fig. 5. Increasing *minPts* and decreasing *ε* have the effect of excluding less dense - or random - cluster formations and breaking up large clusters. For instance in panel (a) at *ε* = 200nm, with *minPts* = 3, 34 clusters are detected. Increasing *minPts* first leads to a decrease of the number of detected clusters (25 clusters for *minPts* = 6), as small, less dense clusters are excluded, and then an increase (30 detected clusters for *minPts* = 30), as larger clusters are broken up. Both effects overlap, and repeat, depending on the statistics of the localizations. In order to exclude the spurious clusters detected at low thresholds, the first steep decrease in cluster densities is excluded by setting *minPts* = 10, see also Fig. S18.

**Figure S18:**
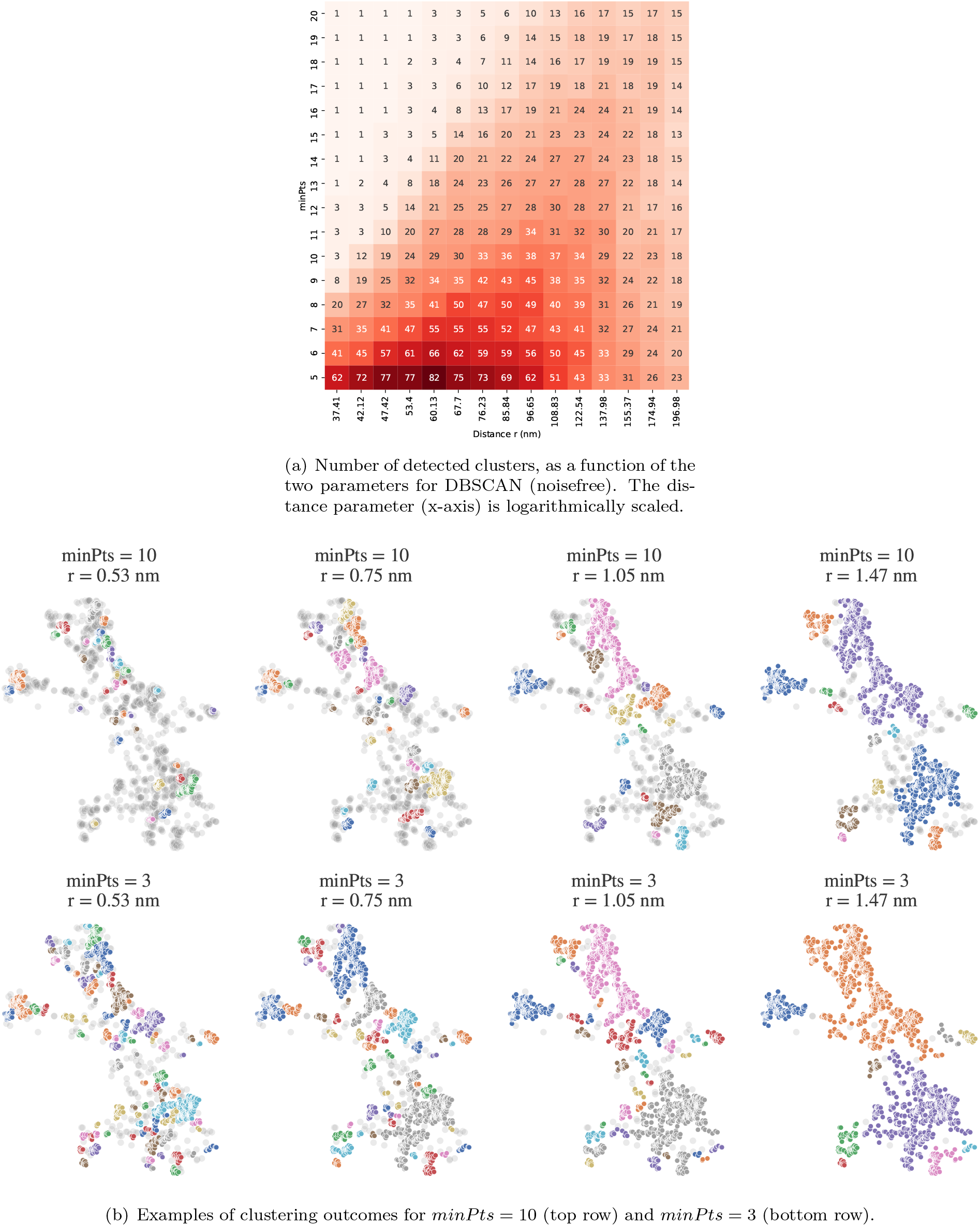
Exploration of clustering outcomes for the localizations in Fig. 5, analysis analogous to Fig. S17.

**Figure S19:**
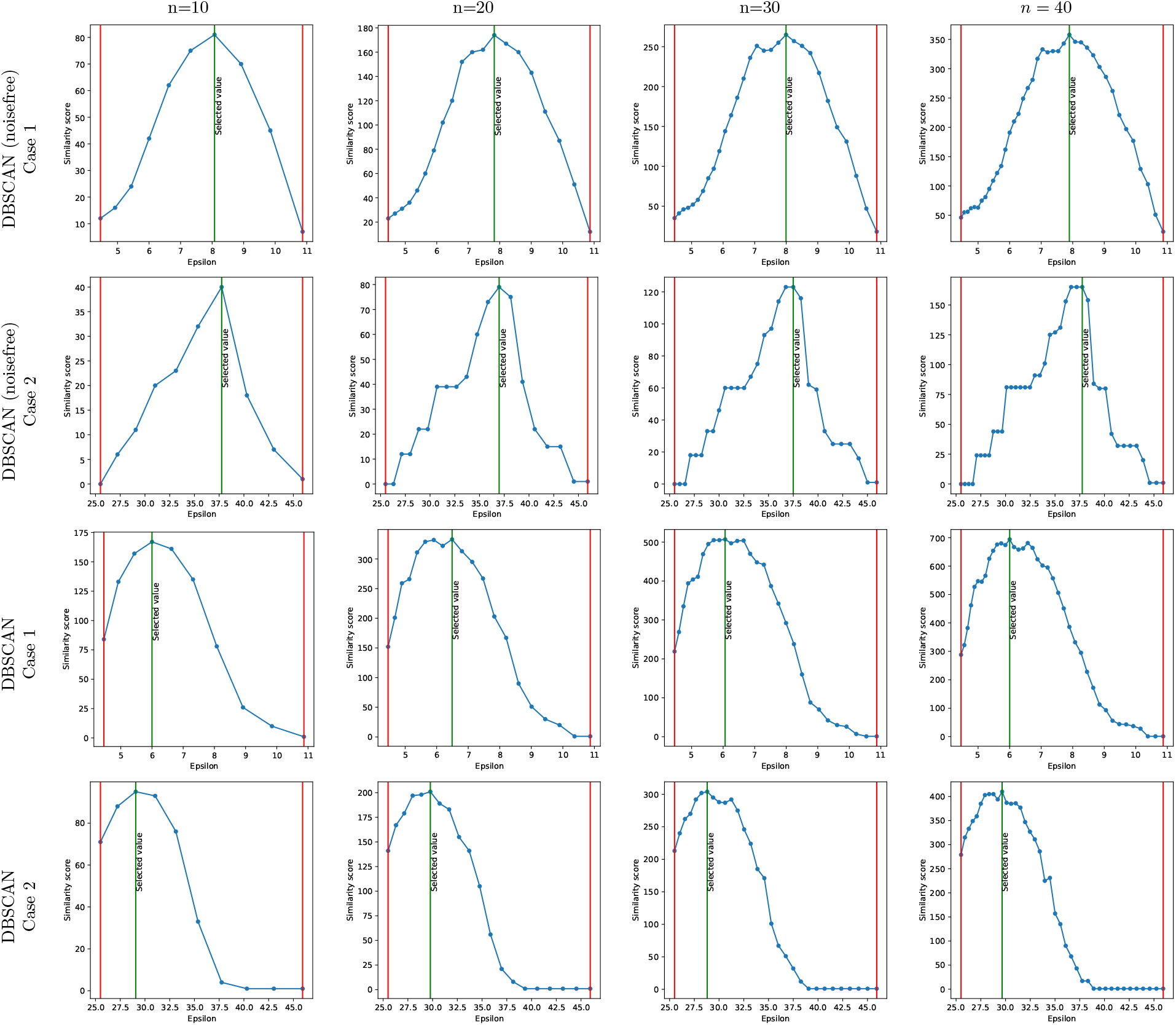
Analysis of examples shown in Figs. S3 and S4, for which the number of *ε*-values within the domain of interest is varied between 10 and 40. It is seen that the similarity score is robust with respect to s in the number of points.

